# AAV-mediated gene therapy produces fertile offspring in the *Lhcgr*-deficient mouse model of Leydig cell failure

**DOI:** 10.1101/2021.04.07.438814

**Authors:** Kai Xia, Fulin Wang, Xingqiang Lai, Peng Luo, Hong Chen, Yuanchen Ma, Weijun Huang, Wangsheng Ou, Yuyan Li, Xin Feng, Zhenmin Lei, Tu Xiang’an, Qiong Ke, Frank F.X. Mao, Chunhua Deng, Andy P. Xiang

## Abstract

Leydig cell failure (LCF) caused by gene mutation results in testosterone deficiency and infertility. Serum testosterone levels can be recovered via testosterone replacement; however, established therapies have shown limited success in restoring fertility. Here, we used a luteinizing hormone/choriogonadotrophin receptor (*Lhcgr*)-deficient mouse model of genetic LCF to investigate the feasibility of gene therapy for restoring testosterone production and fertility. We screened several adeno-associated virus (AAV) serotypes and identified AAV8 as an efficient vector to drive exogenous *Lhcgr* expression in progenitor Leydig cells through interstitial injection. We observed considerable testosterone recovery and Leydig cell maturation after AAV8-Lhcgr treatment in pubertal *Lhcgr*^-/-^ mice. This gene therapy substantially recovered sexual development, partially restored spermatogenesis and effectively produced fertile offspring. Furthermore, these favorable effects could be reproduced in adult *Lhcgr*^-/-^ mice. Our proof-of-concept experiments in this mouse model demonstrate that AAV-mediated gene therapy may represent a promising therapeutic approach for patients with genetic LCF.

## Introduction

Testosterone is the key hormone that regulates the development and maintenance of the masculine phenotype and reproductive function ^1^. Leydig cells (LCs), which are located in the interstitial compartment of the testis and nestled among the seminiferous tubules, are primarily responsible for the production of testosterone via a multistep process involving multiple genes ^2^. When one of these genes is deficient, Leydig cell failure (LCF) occurs; this causes testosterone deficiency, which further contributes to arrested spermatogenesis and infertility ^3, 4^. In such cases, serum testosterone levels can be normalized through testosterone replacement therapy (TRT) ^5^. However, infertility is almost always present in patients with genetic LCF, and conventional established therapies have shown limited success in addressing this issue ^6, 7^. Thus, there is a significant need for new approaches to treat genetic LCF.

Given that the underlying cause of poor testosterone production in LCF is the presence of a genetic variant in a gene involved in regulating steroidogenic pathways ^3^, gene therapy is considered to be one of the most promising potential therapeutic strategies. The replacement of defective genes has been proven to be a suitable therapeutic approach for a number of monogenic diseases in preclinical and clinical studies ^8,9,10,11,12^. Among the available gene delivery vectors, recombinant adeno-associated viruses (AAVs) have become a powerful tool for in vivo gene therapy due to their desirable safety profile ^13^. In the past few years, AAV-mediated gene therapies have been applied to treat many genetic defects using preclinical models, such as those involved in hemophilia ^12^, multiple muscle diseases ^9, 14^, brain disorders ^8^, retinitis pigmentosa ^10^, heart failure ^15^, and deafness ^16^. In addition, several AAV serotypes have been used in clinical trials in patients, including AAV1, AAV2, AAV6, AAV8, AAV9, etc. ^17^. However, whether AAV-mediated gene delivery could be used to restore testosterone production and benefit fertility in genetic LCF remains unknown. Thus, there is an urgent demand for investigating the feasibility of AAV-mediated gene therapy in an animal model of genetic LCF.

Previous reports have described a mouse model of hereditary LCF, which occurs as a result of a null mutation in the gene encoding for *Lhcgr* ^18, 19^. As a highly conserved gene among mammalian species, *Lhcgr* plays important roles in reproduction ^20^. More specifically, *Lhcgr* is required for the maturation of progenitor Leydig cells and the synthesis of testosterone, which regulates the development of sex organs and promotes spermatogenesis ^21^. In *Lhcgr*^-/-^ male mice, the development of Leydig cells is impaired, which leads to a dramatic reduction in testosterone levels. Additionally, *Lhcgr*^-/-^ male mice exhibited stunted sexual development, defective spermatogenesis and infertility ^18, 19^. Mice homozygous for the mutation mimic the phenotype of LCF in humans ^22^ and thus serve as an excellent animal model for gene delivery studies.

Here, we used the *Lhcgr*-deficient mouse model of LCF to first investigate whether AAV-mediated gene therapy can restore Leydig cell function and restart sexual development in pubertal mice. We subsequently elucidated whether this gene therapy could rescue spermatogenesis and the production of fertile offspring. Finally, we investigated the therapeutic effect in adult mice to determine the therapeutic potential of this gene therapy in adult patients who had missed puberty.

## Results

### Testicular injection of AAV8 targets progenitor Leydig cells

Several previous studies suggested that Leydig cell differentiation is blocked in the progenitor cell stage in the postnatal testis of *Lhcgr*-deficient LCF mice ^21, 23^. Because different AAV serotypes have shown distinct tissue or cell tropisms, to identify the AAV serotypes with the highest viral transduction rate towards progenitor Leydig cells, we interstitially injected AAV-CAG-mCherry reporter vectors containing the commonly used AAV serotypes 1, 2, 6, 8, or 9 at titers of 8×10^10 genome copies (gc) into each testis of *Lhcgr*^-/-^ mice. We performed histological analysis of the testes at 7 days after vector injection and found that, at equivalent viral titers, AAV8 exhibited the highest efficiency among the tested serotypes as demonstrated by co-expression of mCherry and progenitor Leydig cells markers Nestin and platelet-derived growth factor receptor α (Pdgfrα) (Fig. S1A, B). AAV8 was the most effective serotype among all the tested AAV, which successfully transduced of over 80% progenitor Leydig cells on average in the interstitium of the testis (Fig. S1C, D). Thereafter, we evaluated the tropism of AAV8 for germ cells and Sertoli cells. Immunostaining analysis revealed that mCherry was not co-expressed with the germ cell marker DEAD-box helicase 4 (Ddx4) or the Sertoli cell marker SRY-box transcription factor 9 (Sox9), indicating the absence of AAV infection in these two cell types (Fig. S1E, F). Our results indicate that the interstitial injection of AAV8 could effectively and exclusively target progenitor Leydig cells in *Lhcgr*^-/-^ mice.

### Gene delivery of AAV8-Lhcgr increases *Lhcgr* expression in the testes and recovers testosterone levels in pubertal *Lhcgr*^-/-^ mice

Next, we used an established *Lhcgr*^-/-^ mouse model of LCF to determine whether AAV-based gene therapy could be efficacious in recovering *Lhcgr* expression and testosterone levels. We generated AAV8 vectors that carried the coding sequence of mouse *Lhcgr* with the CAG promoter (AAV8-Lhcgr), and these vectors were used in subsequent experiments (Fig. 1A). To evaluate the functionality of these vectors, we first chose pubertal *Lhcgr*^-/-^ mice (3 weeks old) to determine the potential therapeutic effects of this gene therapy in LCF patients at puberty (Fig. 1B). These mice were interstitially injected with phosphate-buffered saline (PBS) or AAV8-Lhcgr at doses of 8×10^9, 4×10^10, 8×10^10 or 2×10^11 gc/testis. Untreated littermate *Lhcgr*^+/+^ and *Lhcgr*^+/-^ mice served as controls. To evaluate the efficiency of gene transfer, we measured RNA transcript and relative protein levels at 4 weeks after AAV8-Lhcgr administration. Quantitative RT-PCR analysis of testicular tissue showed that the administration of AAV8-Lhcgr resulted in dose-dependent expression of *Lhcgr* transcripts in testes from injected *Lhcgr*^-/-^ mice, whereas *Lhcgr* expression was not detectable in *Lhcgr*^-/-^ mice treated with PBS (Fig. 1C). Accordingly, immunostaining demonstrated obvious *Lhcgr* expression in the testicular interstitium in the AAV8-Lhcgr-treated group (8×10^10 gc/testis), whereas *Lhcgr* was negligibly detected in the testicular interstitium from *Lhcgr*^-/-^ mice treated with PBS (Fig. S2).

**Fig. 1.**
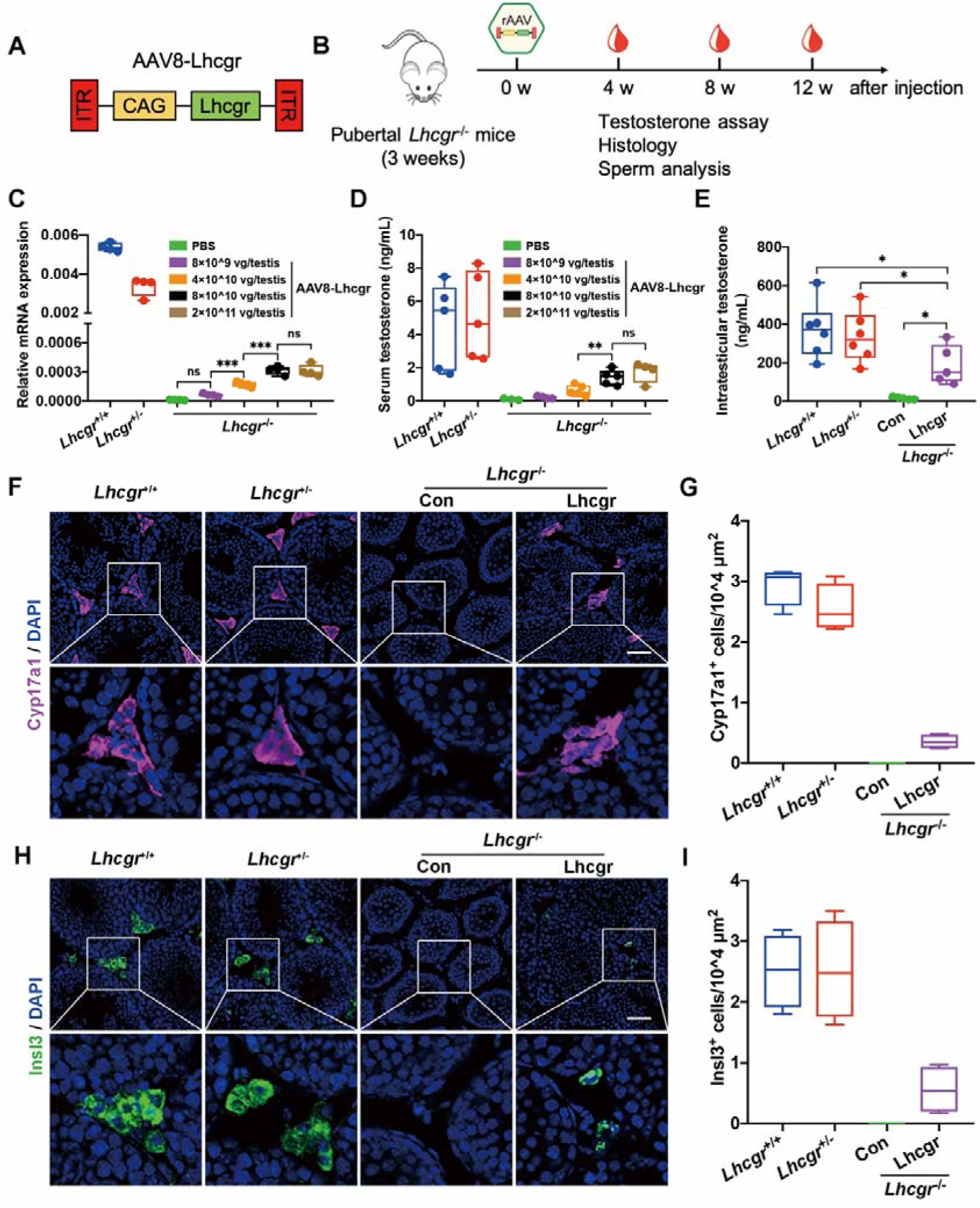
Testicular injection of AAV8-Lhcgr rescues Leydig cell function and recovers testosterone levels in pubertal *Lhcgr*^-/-^ mice. (A) Schematic of the AAV vector used in the study. (B) Experimental overview of the in vivo studies. (C) Quantitative RT-PCR was used to quantify *Lhcgr* mRNA transcripts in testicular tissues from *Lhcgr*^+/+^ mice (n=4), *Lhcgr*^+/-^ mice (n=4), and *Lhcgr*^-/-^ mice injected with PBS (n=4) or increasing doses of AAV8-Lhcgr [8×10^9 gc/testis (n=4), 4×10^10 gc/testis (n=4), 8×10^10 gc/testis (n=4), and 2×10^11 gc/testis (n = 4)] at 4 weeks after treatment. β-actin was used for normalization. (D) The concentrations of serum testosterone were analyzed 4 weeks after treatment in *Lhcgr*^+/+^ mice (n=5), *Lhcgr*^+/-^ mice (n=5), and *Lhcgr*^-/-^ mice injected with PBS (n=3) or increasing doses of AAV8-Lhcgr [8×10^9 gc/testis (n=4), 4×10^10 gc/testis (n=5), 8×10^10 gc/testis (n=5), and 2×10^11 gc/testis (n=4)]. (E) The concentrations of intratesticular testosterone were detected 4 weeks after treatment in *Lhcgr*^+/+^ mice (n=6), *Lhcgr*^+/-^ mice (n=6), and *Lhcgr*^-/-^ mice injected with PBS (n=5) or AAV8-Lhcgr (8×10^10 gc/testis, n=5). (F) The Leydig cell marker, Cyp17a1, in the testicular interstitium was evaluated by immunofluorescence assay at 4 weeks after treatment in *Lhcgr*^+/+^ mice (n=4), *Lhcgr*^+/-^ mice (n=4), and *Lhcgr*^-/-^ mice injected with PBS (n=4) or AAV8-Lhcgr (8×10^10 gc/testis, n=4). Scale bar: 100 μm. (G) Cyp17a1^+^ cells were quantified in the different groups (n=4 per group). (H) Immunostaining with anti-Insl3 antibody was used to detect mature Leydig cells at 4 weeks after treatment in *Lhcgr*^+/+^ mice (n=4), *Lhcgr*^+/-^ mice (n=4), and *Lhcgr*^-/-^ mice injected with PBS (n=4) or AAV8-Lhcgr (8×10^10 gc/testis, n=4). Scale bar: 100 μm. (I) Quantification of Insl3^+^ cells was performed in the different groups (n=4 per group). Data are represented by box plots, and whiskers are minimum to maximum values. Significance was determined by one-way ANOVA. * P < 0.05, ** P < 0.01, *** P < 0.001, ns = not significant.

In addition, serum testosterone concentrations were significantly and dose-dependently increased in *Lhcgr*^-/-^ mice treated with AAV8-Lhcgr compared with PBS. Serum testosterone levels in mice treated with 8×10^10 and 2×10^11 gc/testis AAV8-Lhcgr reached approximately 30% the levels observed in *Lhcgr*^+/+^ or *Lhcgr*^+/-^ mice, whereas the level was profoundly lower (nearly undetectable) in PBS-treated *Lhcgr*^-/-^ males, suggesting that *Lhcgr*^-/-^ mice exhibited recovery of testosterone production after receiving AAV8-Lhcgr (Fig. 1D). Significant restoration of serum testosterone levels was also detected in *Lhcgr*^-/-^ mice at 8 and 12 weeks after treatment with AAV8-Lhcgr at the dose of 8×10^10 gc/testis (Fig. S3A, B). Notably, the concentration of intratesticular testosterone, which is considered to be vital for spermatogenesis ^7^, was also increased in AAV8-Lhcgr-treated mice (8×10^10 gc/testis) compared with those in the PBS-treated group at 4 weeks after treatment (Fig. 1E).

### AAV8-Lhcgr promotes Leydig cell maturation in pubertal *Lhcgr*^-/-^ mice

We next used molecular analyses to assess the impact of AAV8-Lhcgr on Leydig cell maturation in the pubertal cohort. Four weeks after AAV8-Lhcgr injection, we examined the expression levels of Leydig cell markers in testes from *Lhcgr*^+/+^, *Lhcgr*^+/-^, and *Lhcgr*^-/-^ mice injected with PBS or AAV8-Lhcgr (8×10^10 gc/testis). Immunofluorescence analysis showed increased Leydig cell numbers in AAV-treated *Lhcgr*^-/-^ mice compared to PBS-treated mice as determined by staining of cytochrome P450 family 17 subfamily A member 1 (Cyp17a1) in the interstitial space of testes samples (Fig. 1F, G). Furthermore, AAV8-Lhcg treatment restored the maturation of Leydig cells to approximately 22% of that observed in *Lhcgr*^+/+^ or *Lhcgr*^+/-^ mice as determined by insulin-like peptide 3 (Insl3) staining in the testicular interstitium, whereas this marker was only negligibly detected in the testicular interstitium of PBS-treated *Lhcgr*^-/-^ mice (Fig. 1H, I). These data suggest that AAV8-Lhcgr treatment in pubertal *Lhcgr*^-/-^ mice partially restored the level of mature Leydig cells.

### AAV8-Lhcgr restarts sexual development in pubertal *Lhcgr*^-/-^ mice

Based on the observation that testosterone levels and the maturation of Leydig cells from *Lhcgr*^-/-^ mice were promoted after AAV8-Lhcgr therapy (8×10^10 gc/testis), we next examined whether these features were accompanied by normalization of sexual development. Intriguingly, in *Lhcgr*^-/-^ mice treated with AAV8-Lhcgr, we found that the retained testes descended into the scrotum at 4 weeks after gene delivery. Moreover, the underdeveloped external genitals (penis and scrotum) of the *Lhcgr*^-/-^ mice approached the size of those observed in *Lhcgr*^+/+^ and *Lhcgr*^+/-^ mice (Fig. 2A, B).

**Fig. 2.**
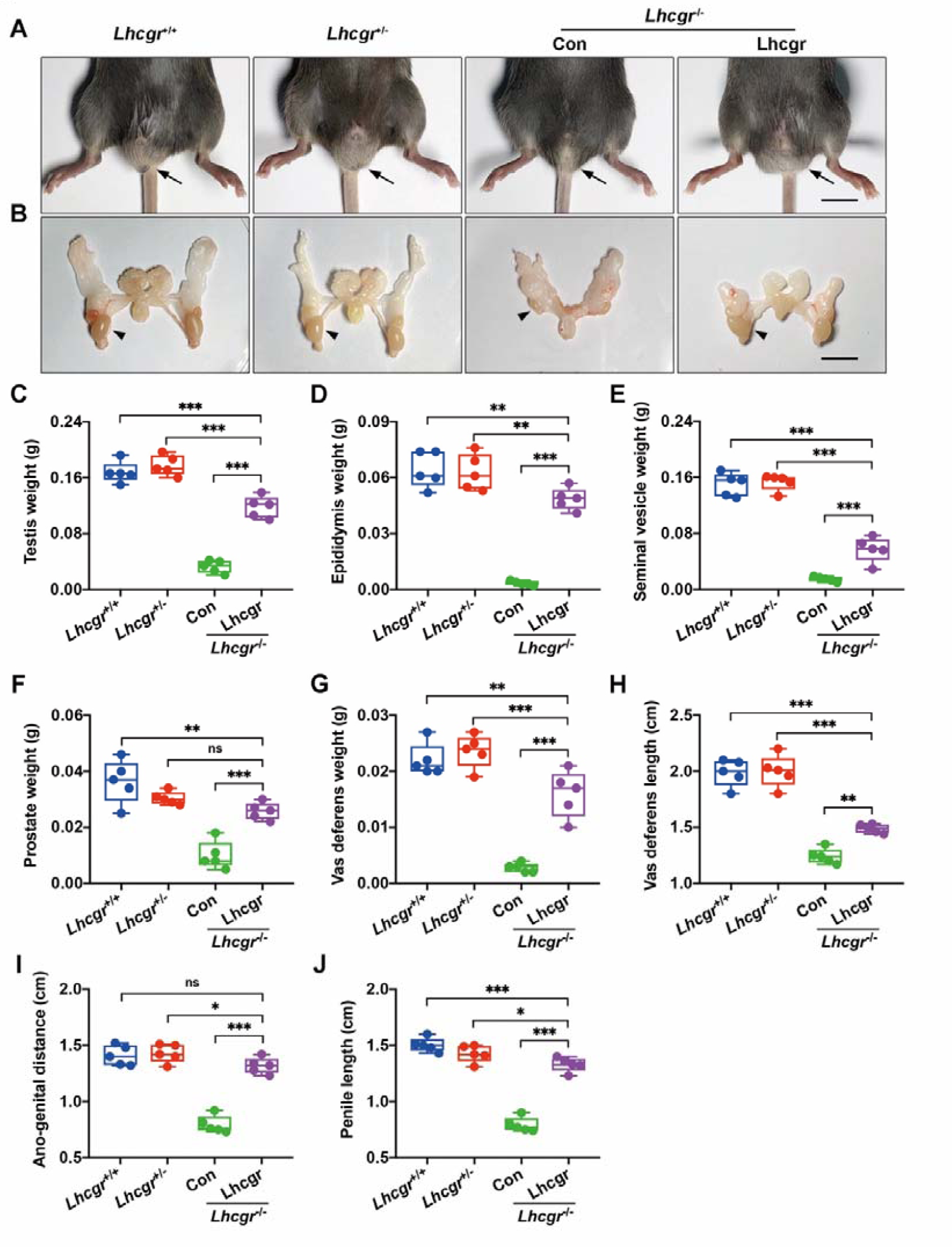
AAV8-Lhcgr restarts sexual development in pubertal *Lhcgr*^-/-^ mice. (A,B) Representative photographs of external (A) and internal genitalia (B) of *Lhcgr*^+/+^ mice (n=5), *Lhcgr*^+/-^ mice (n=5), and *Lhcgr*^-/-^ mice injected with PBS (n=5) or AAV8-Lhcgr (8×10^10 gc/testis, n=5) at 4 weeks after treatment. Arrows (A) and arrowheads (B) indicate the testes. Scale bar: 1 cm. (C-J) Testis weight (C), epididymis weight (D), seminal vesicle weight (E), prostate weight (F), vas deferens weight (G), vas deferens length (H), ano-genital distance (I), and penile length (J) of *Lhcgr*^+/+^ mice (n=5), *Lhcgr*^+/-^ mice (n=5), and *Lhcgr*^-/-^ mice injected with PBS (n=5) or AAV8-Lhcgr (8×10^10 gc/testis, n=5) at 4 weeks after treatment. Data are represented by box plots, and whiskers are minimum to maximum values. Significance was determined by one-way ANOVA. * P < 0.05, ** P < 0.01, *** P < 0.001, ns = not significant.

Further examination revealed that the testis weights of AAV8-Lhcgr-treated mice were approaching to the level of those in *Lhcgr*^+/+^ or *Lhcgr*^+/-^ mice, whereas testis weights in the PBS-treated group remained far lower (Fig. 2C). In addition, the hypoplastic epididymis of *Lhcgr*^-/-^ mice grew markedly under AAV8-Lhcgr treatment, almost reaching the size observed in *Lhcgr*^+/+^ and *Lhcgr*^+/-^ mice (Fig. 2D). The seminal vesicle weight, which is a well-established biomarker of androgen exposure ^24^, increased in the AAV8-Lhcgr group to approximately 37% of the weights in *Lhcgr*^+/+^ and *Lhcgr*^+/-^ mice, whereas the seminal vesicles were macroscopically undetectable in PBS-treated *Lhcgr*^-/-^ mice (Fig. 2E). The prostates of AAV8-Lhcgr-treated Lhcgr^-/-^ mice weighed more than those of the PBS-treated group and were more than half of the prostate weight in *Lhcgr*^+/+^ and *Lhcgr*^+/-^ mice (Fig. 2F). The weight and length of the vas deferens were higher in AAV8-Lhcgr-treated *Lhcgr*^-/-^ mice than in PBS-treated group, although these parameters remained significantly lower than those recorded in in the *Lhcgr*^+/+^ and *Lhcgr*^+/-^ groups (Fig. 2G, H). The ano-genital distances were increased in AAV8-Lhcgr-treated *Lhcgr*^-/-^ mice compared with the PBS-treated group (Fig. 2I), indicating that masculinization was promoted after gene therapy. AAV8-Lhcgr treatment also increased penile length compared with that of PBS-treated *Lhcgr*^-/-^ mice (Fig. 2J). Overall, these results strongly support the notion that AAV8-Lhcgr restarts sexual development in pubertal *Lhcgr*^-/-^ mice.

### AAV8-Lhcgr rescues spermatogenesis in pubertal *Lhcgr*^-/-^ mice

Given that AAV8-Lhcgr treatment could recover testosterone levels and promote sexual development in pubertal *Lhcgr*^-/-^ mice, we next investigated whether AAV8-Lhcgr could rescue spermatogenesis. Histological analysis of the PBS-injected *Lhcgr*^-/-^ testes showed that the seminiferous tubules were decreased in size and that spermatogenesis was arrested without any mature spermatids, as reported previously ^18, 19^. In *Lhcgr*^-/-^ mice treated with AAV8-Lhcgr (8×10^10 gc/testis), the width of the seminiferous tubules was increased, and spermatogenesis was evident in these testes (Fig. 3A). Quantification of morphological changes revealed that the seminiferous tubule diameter was 80.6±15.0 μm in the testes of PBS-treated *Lhcgr*^-/-^ mice and increased to 150.5±21.6 μm in the AAV8-Lhcgr group, approaching levels observed in *Lhcgr*^+/+^ (162.7±22.7 μm) and *Lhcgr*^+/-^ (161.6±21.3 μm) tubules (Fig. 3B). When calculating the percentage of seminiferous tubules that contained mature spermatids in the testes, we found that approximately one-third of the tubules in *Lhcgr*^-/-^ mice injected with AAV8-Lhcgr contained mature spermatids, indicating that full spermatogenesis occurred in this group (Fig. 3C).

**Fig. 3.**
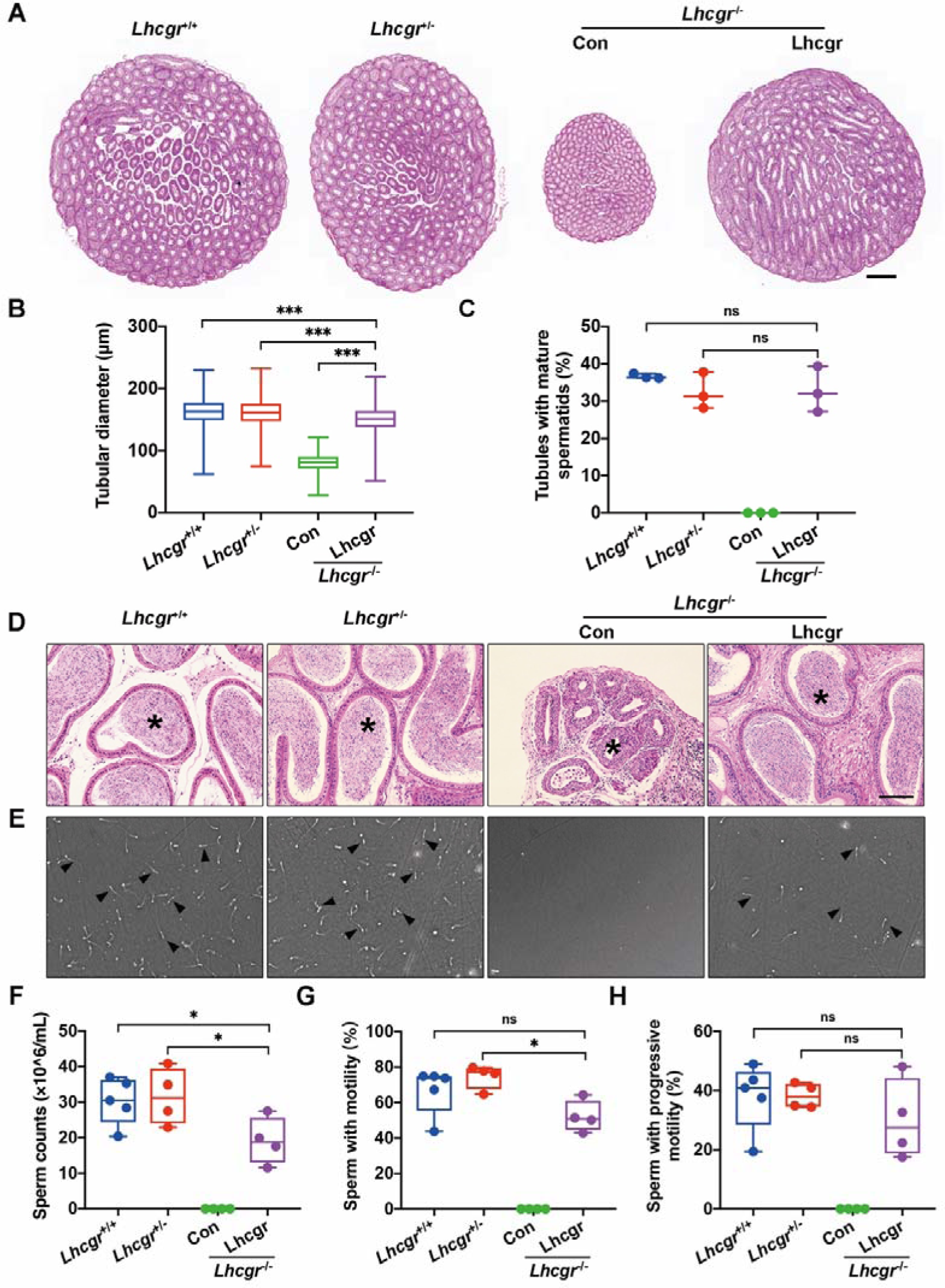
AAV8-Lhcgr rescues spermatogenesis. (A) Representative light micrographs of testis sections from *Lhcgr*^+/+^ mice (n=3), *Lhcgr*^+/-^ mice (n=3), and *Lhcgr*^-/-^ mice injected with PBS (n=3) or AAV8-Lhcgr (8×10^10 gc/testis, n=3). Samples were taken at 7 weeks of age (4 weeks after treatment). Scale bars: 500 μm. (B,C) The diameter of seminiferous tubules (B) and the percentage of tubules with mature spermatids (C) were counted in the testis of *Lhcgr*^+/+^ mice (256-281 tubules calculated per specimen, n=3), *Lhcgr*^+/-^ mice (256-288 tubules calculated per specimen, n=3), and *Lhcgr*^-/-^ mice injected with PBS (251-259 tubules calculated per specimen, n=3) or AAV8-Lhcgr (8×10^10 gc/testis, 188-276 tubules calculated per specimen, n=3). (D) Histological analysis of epididymis collected from *Lhcgr*^+/+^ mice (n=3), *Lhcgr*^+/-^ mice (n=3), and *Lhcgr*^-/-^ mice injected with PBS (n=3) or AAV8-Lhcgr (8×10^10 gc/testis, n=3). Scale bars: 100 μm. (E) Representative light micrographs of sperm obtained from the cauda epididymis of *Lhcgr*^+/+^ mice (n=5), *Lhcgr*^+/-^ mice (n=4), and *Lhcgr*^-/-^ mice injected with PBS (n=4) or AAV8-Lhcgr (8×10^10 gc/testis, n=4). Scale bars: 100 μm. (F-H) The sperm concentration and proportions of sperm with motility and progressive motility were analyzed 4 weeks after treatment [*Lhcgr*^+/+^ mice (n=5), *Lhcgr*^+/-^ mice (n=4), and *Lhcgr*^-/-^ mice injected with PBS (n=4) or AAV8-Lhcgr (8×10^10 gc/testis, n=4)]. Data are represented by box plots, and whiskers are minimum to maximum values. Significance was determined by Kruskal-Wallis test (B) or one-way ANOVA (C, F-H). * P < 0.05, ** P < 0.01, *** P < 0.001, ns = not significant.

To further characterize spermatogenesis after gene therapy, epididymis samples were collected from the four groups at 4 weeks after treatment. Histological analysis of the epididymis showed that the luminal diameters of tubules in the cauda were dramatically decreased in *Lhcgr*^-/-^ mice compared to *Lhcgr*^+/+^ or *Lhcgr*^+/-^ group, and the lumens of the former were completely devoid of spermatids. Treatment of *Lhcgr*^-/-^ mice with AAV8-Lhcgr (8×10^10 gc/testis) restored the luminal diameter in the cauda epididymis and was associated with the presence of many spermatids at this location (Fig. 3D). To further quantify the degree of spermatogenesis observed after AAV8-Lhcgr treatment, the quantity and motility of sperm were determined using a computer-aided semen analysis (CASA) system (Fig. 3E). At 4 weeks post-treatment, the epididymal sperm number of the AAV8-Lhcgr group was increased to over half of that observed in *Lhcgr*^+/+^ and *Lhcgr*^+/-^ mice, whereas sperm were not detected in PBS-treated *Lhcgr*^-/-^ mice (Fig. 3F). Analysis of sperm progressive motility at 4 weeks post-treatment revealed no apparent difference in the sperm of AAV8-Lhcgr-treated *Lhcgr*^-/-^ mice and their *Lhcgr*^+/+^ or *Lhcgr*^+/-^ littermates (Fig. 3G, H; Video S1, 2). Collectively, these results suggest that AAV8-Lhcgr rescues spermatogenesis and qualitatively increases sperm number and motility.

### AAV8-Lhcgr promotes the formation of round and elongating spermatids

To molecularly define the consequences of AAV8-Lhcgr on spermatogenesis, RNA sequencing (RNA-seq) was performed at 4 weeks after AAV8-Lhcgr (8×10^10 gc/testis) treatment. RNA-seq analysis showed that AAV8-Lhcgr-injected testes of *Lhcgr*^-/-^ mice had an extremely high similarity in gene expression with *Lhcgr*^+/+^ and *Lhcgr*^+/-^ groups, whereas the PBS-treated *Lhcgr*^-/-^ group showed less similarity with the other three groups (Fig. S4A, B). Furthermore, we performed Gene Ontology (GO) analysis of the differentially expressed genes (DEGs) between PBS and AAV8-Lhcgr-treated *Lhcgr*^-/-^ testes (Fig. 4A). The results showed that upregulated genes were enriched for “germ cell development” and “spermatid differentiation”, indicating that processes involved in spermatogenesis were activated after AAV8-Lhcgr treatment. To determine the stages of spermatogenesis at which AAV8-Lhcgr gene therapy functions, we queried these data with functionally defined genes reflecting spermatogonia, spermatocytes, round spermatocytes, and elongating spermatids ^25, 26^ and observed that the transcript profile of *Lhcgr*^-/-^ testis treated with PBS was enriched for genes related to spermatogonia (Dazl, Stra8, Zbtb16, etc.) and spermatocytes (Tex101, Piwil1, Sycp3, etc.). However, AAV8-Lhcgr-treated samples were highly enriched in transcripts specific for round spermatids (Acrv1, Tssk1, Spag6, Spaca1, etc.) and elongating spermatids (Best1, Pabpc1, Ccdc89, Prm1, etc.) (Fig. 4B). These results were further confirmed by quantitative RT-PCR analysis (Fig. S5A-D). Immunofluorescence analysis of peanut agglutinin (PNA), which labels the acrosome, supported the appearance of spermatids in the testes of AAV8-Lhcgr-treated *Lhcgr*^-/-^ mice, whereas PNA^+^ signals were incredibly weak in the PBS-treated group (Fig. 4C, D). Transition protein 2 (Tnp2), which is expressed in the nuclei of elongating spermatids during histone-to-protamine transition ^27^, was barely detected in the testes of PBS-treated *Lhcgr*^-/-^ mice, whereas its expression was clearly detected in those of AAV8-Lhcgr-treated mice (Fig. 4E, F). Together, these findings support the hypothesis that AAV8-Lhcgr treatment promotes the formation of round and elongating spermatids.

**Fig. 4.**
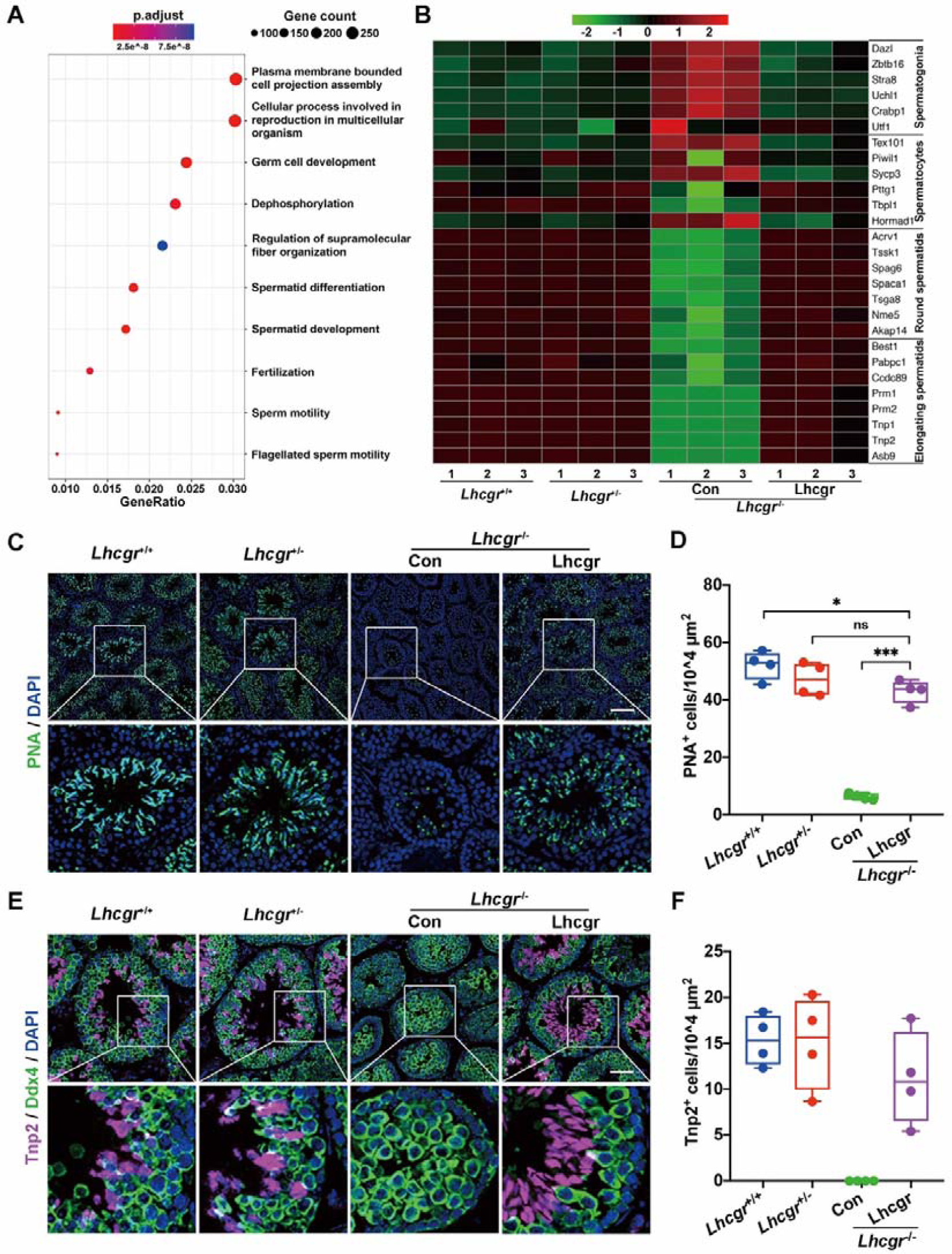
AAV8-Lhcgr promotes the formation of elongating spermatids. (A) Representative GO terms of the top 500 upregulated genes in male *Lhcgr*^-/-^ mice treated with AAV8-Lhcgr (8×10^10 gc/testis, n=3) versus PBS (n=3). (B) Heatmap showing expression of marker genes for the four major germ cell types in the testicular transcriptional profiles of *Lhcgr*^+/+^ mice (n=3), *Lhcgr*^+/-^ mice (n=3), and *Lhcgr*^-/-^ mice injected with PBS (n=3) or AAV8-Lhcgr (8×10^10 gc/testis, n=3). Shown are the mean expression levels for marker genes of each germ cell type. Representative markers are listed on the right side. (C-F) Representative images of testis sections from *Lhcgr*^+/+^ mice (n=4), *Lhcgr*^+/-^ mice (n=4), and *Lhcgr*^-/-^ mice injected with PBS (n=4) or AAV8-Lhcgr (8×10^10 gc/testis, n=4). Sections were immunostained for PNA (C), Ddx4 (a marker of germ cells), and Tnp2 (E), and counterstained with DAPI. Quantitative analysis showing the percentage of PNA^+^ (D) and Tnp2^+^ (F) germ cells in the seminiferous tubules of the testes. Scale bars: 50 μ Data are represented by box plots, and whiskers are minimum to maximum values. Significance was determined by one-way ANOVA. * P < 0.05, ** P < 0.01, *** P < 0.001.

### AAV8-Lhcgr restores fertility and produces fertile offspring

To assess whether the sperm produced after gene therapy could support reproduction, in vitro fertilization (IVF) was performed using spermatids obtained from the caudal epididymis of male *Lhcgr*^-/-^ mice at 4 weeks after AAV8-Lhcgr (8×10^10 gc/testis) injection and oocytes harvested from female *Lhcgr*^+/+^ mice (Fig. 5A). Among a total of 723 eggs used for IVF, 178 eggs (24.6%) successfully progressed to 2-cell embryos in vitro (efficiency, 12.5% to 42.4%). Of these 2-cell embryos, 149 were transplanted into the uterus of 8 pseudo-pregnant mice (Fig. 5B; Table S1), which produced 58 offspring (efficiency 9.6% to 55.6%) (Fig. 5C; Table S1). To confirm that the offspring were derived from the AAV8-Lhcgr-treated *Lhcgr*^-/-^ male mice and *Lhcgr*^+/+^ females, eight pups were subjected to PCR-based genotyping (Fig. 5D). The results showed that the eight pups carried the wild-type and mutated alleles at proportions consistent with Mendelian law (Fig. 5D).

**Fig. 5.**
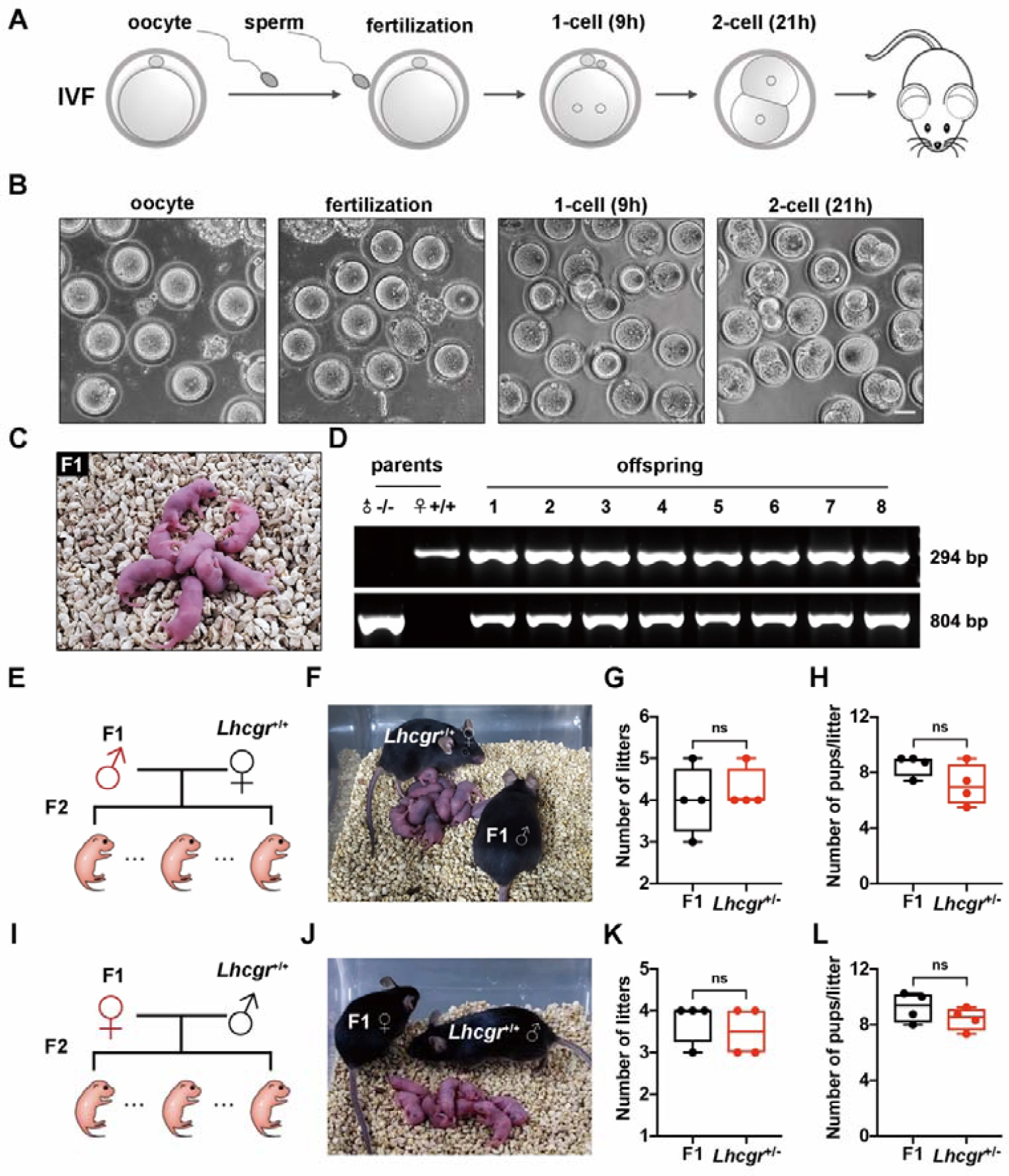
AAV8-Lhcgr restores fertility and produces fertile offspring. (A) The in vitro fertilization (IVF) scheme used to produce offspring with sperm from AAV8-Lhcgr-treated *Lhcgr*^-/-^ mice (8×10^10 gc/testis, n=2) and oocytes from *Lhcgr*^+/+^ females. (B) Representative images of oocytes, fertilization, 1-cell embryo, and 2-cell embryo. Scale bar: 50 μm. (C) The offspring (F1) derived from AAV8-Lhcgr-treated *Lhcgr*^-/-^ male mice. (D) Genotyping of the pups derived from AAV8-Lhcgr-treated *Lhcgr*^-/-^ males and *Lhcgr*^+/+^ females. The amplified wild-type (wt) DNA is 294 bp while the mutant (mt) DNA fragment is 804 bp. (E) Mating scheme used to produce the second generation (F2) mice. (F) The male F1 mice were used to produce F2 by mating with *Lhcgr*^+/+^ females. (G,H) Continuous breeding assay starting at 6 weeks of age, showing numbers of litters (G) and pups per litter (H) between F1 males (n=4) and *Lhcgr*^+/-^ males (n=4) within 4 months. (I) Mating scheme used to produce F2 mice. (J) The female F1 mice were used to produce F2 pups by mating with *Lhcgr*^+/+^ males. (K,L) Continuous breeding assay starting at 6 weeks of age, showing numbers of litters (K) and pups per litter (L) between F1 females (n=4) and *Lhcgr*^+/-^ females (n=4) within 4 months. Data are represented by box plots, and whiskers are minimum to maximum values. Significance was determined by Student’s t-test. ns = not significant.

To test whether AAV8 was integrated into the genomes of offspring, we performed PCR using vector-specific primers for the CAG promoter and the inserted *Lhcgr*. We analyzed tail DNA of eight representative offspring born from the AAV8-Lhcgr transduction experiments but failed to detect any vector sequences (Fig. S6A, B). This finding indicates that AAV8 did not undergo integration into the genomes of offspring. We next asked whether the offspring created via AAV8-Lhcgr gene therapy could produce a second generation. Four mature males and four females generated from AAV8-Lhcgr-treated *Lhcgr*^-/-^ mice were mated to corresponding *Lhcgr*^+/+^ mice, and all were proven to be fertile (Fig. 5E, F, I, J). Moreover, the offspring from AAV8-Lhcgr-treated *Lhcgr*^-/-^ mice exhibited normal fertility compared to that of *Lhcgr*^+/-^ mice (Fig. 5G, H, K, L). Collectively, AAV8-Lhcgr treatment in pubertal *Lhcgr*^-/-^ mice gives rise to fertile offspring.

### AAV8-Lhcgr recovers testosterone levels and rescues spermatogenesis in adult *Lhcgr*^-/-^ mice

Because adult LCF patients miss the optimal opportunity for treatment at puberty ^28,29, 30^, we next question whether this approach still has therapeutic potential in adult mice. Eight-week-old *Lhcgr*^-/-^ mice were injected with AAV8-Lhcgr (8×10^10 gc/testis) and the effects were evaluated 4 weeks later (Fig. 6A). Similar to the results obtained in the pubertal cohort, we found that administration of AAV8-Lhcgr to these *Lhcgr*^-/-^ mice increased *Lhcgr* transcript expression in testes compared with that in the PBS group (Fig. 6B). In parallel with increased *Lhcgr* expression, serum and intratesticular testosterone levels were recovered in AAV8-Lhcgr-injected *Lhcgr*^-/-^ mice (Fig. 6C, D). Immunofluorescence assays revealed the expression of Leydig cell markers Cyp17a1 and Insl3 (Fig. S7A-D), suggesting that Leydig cell maturation occurred in AAV8-Lhcgr-treated *Lhcgr*^-/-^ mice. We also observed normalization of sexual development in *Lhcgr*^-/-^ mice after AAV8-Lhcgr treatment (Fig. 6E, F; S8A-H). Notably, AAV8-Lhcgr-treated mice achieved significant recovery of spermatogenesis as determined by morphological changes of the testis (Fig. 6G), the presence of sperm in the caudal epididymis (Fig. 6H), and significantly increased semen parameters (Fig. 6I-K). The recovery of spermatogenesis was also molecularly characterized by quantitative RT-PCR for round and elongating spermatid-specific genes and immunostaining for PNA and Tnp2. These parameters were upregulated in AAV8-Lhcgr-treated *Lhcgr*^-/-^ testes compared with PBS-treated testes (Fig. S9A-F).

**Fig. 6.**
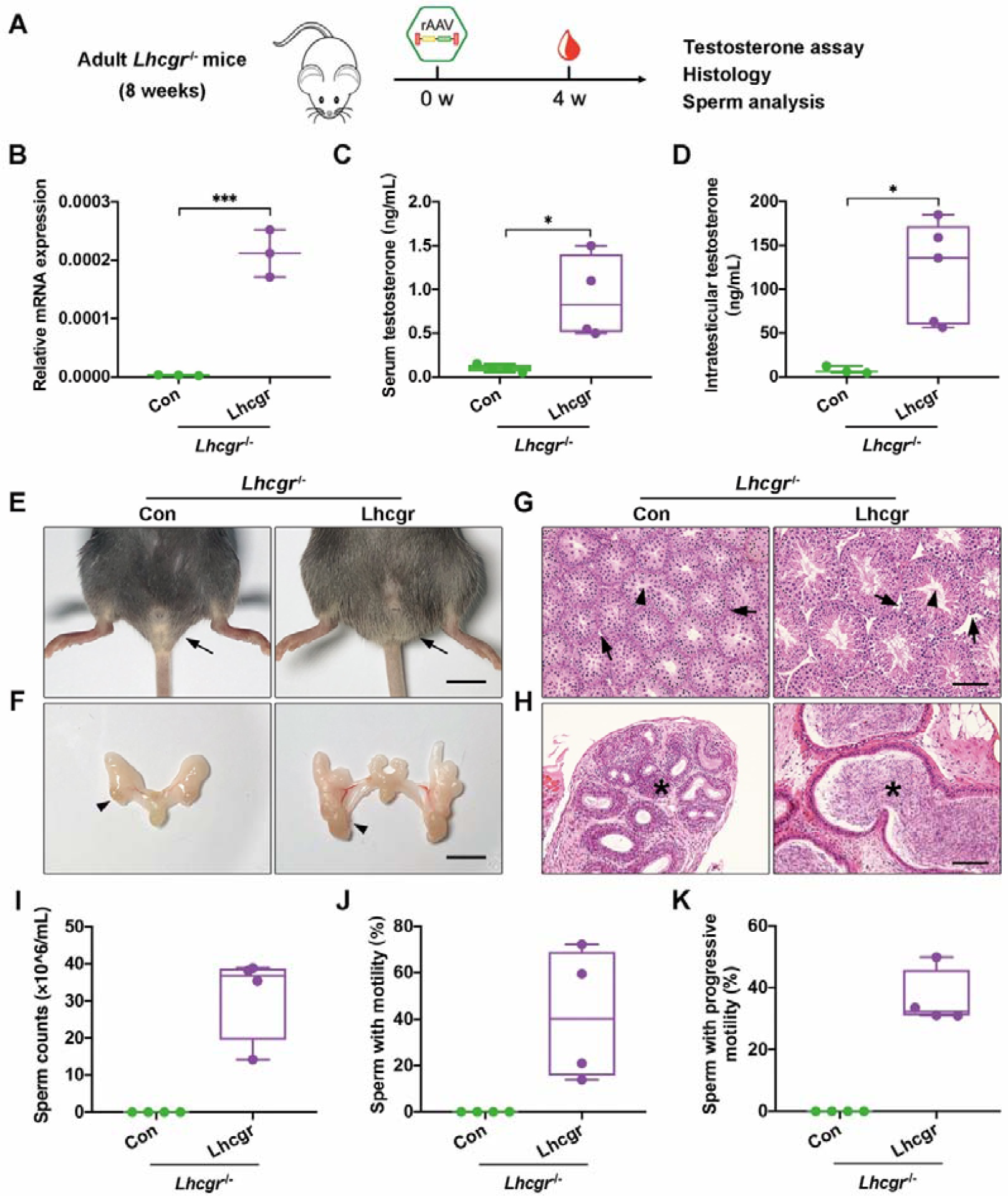
AAV8-Lhcgr restarts sexual development and rescues spermatogenesis in adult *Lhcgr*-deficient mice. (A) Experimental overview of the in vivo studies. (B) Quantitative RT-PCR was used to quantify *Lhcgr* mRNA transcripts in testis tissues from *Lhcgr*^-/-^ mice injected with PBS (n=3) or AAV8-Lhcgr (8×10^10 gc/testis, n=3) at 4 weeks after treatment. β-actin was used for normalization. (C) The concentrations of serum testosterone were analyzed in *Lhcgr*^-/-^ mice injected with PBS (n=4) or AAV8-Lhcgr (8×10^10 gc/testis, n=4) at 4 weeks after treatment. (D) The concentrations of intratesticular testosterone were analyzed 4 weeks after treatment in *Lhcgr*^-/-^ mice injected with PBS (n=3) or AAV8-Lhcgr (8×10^10 gc/testis, n=5). (E,F) Representative photographs of external and internal genitalia of *Lhcgr*^-/-^ mice injected with PBS (n=4) or AAV8-Lhcgr (8×10^10 gc/testis, n=5) at 4 weeks after treatment. Arrows (E) and arrowheads (F) indicate the testes. Scale bar: 1 cm. (G,H) Representative light micrographs of testis sections (G) or cauda epididymis (H) from *Lhcgr*^-/-^ mice injected with PBS (n=4) or AAV8-Lhcgr (8×10^10 gc/testis, n=5). Samples were taken at 12 weeks of age (4 weeks after treatment). Arrows indicate Leydig cells and arrowheads indicate full spermatogenesis in testis (G). Stars indicates spermatozoa in the cauda epididymis (H). Scale bars: 100 μm. (I-K) The sperm concentration and proportions of sperm with motility and progressive motility were analyzed 4 weeks after treatment [*Lhcgr*^-/-^ mice injected with PBS (n=4) or AAV8-Lhcgr (8×10^10 gc/testis, n=4)]. Data are represented by box plots, and whiskers are minimum to maximum values. Significance was determined by Student’s t-test. * P < 0.05, ** P < 0.01, *** P < 0.001, ns = not significant.

To further investigate the time window of AAV8-Lhcgr gene therapy, 6-month-old mice were enrolled in the following experiments (Fig. S10A). Consistent with our findings in the 3- and 8-week-old cohorts, we observed improvements in testosterone levels (Fig. S10B, C) and Leydig cell maturation (Fig. S10D) in AAV8-Lhcgr-treated *Lhcgr*^-/-^ mice (8×10^10 gc/testis). Moreover, AAV8-Lhcgr promoted spermatogenesis in these mice as inferred by the appearance of spermatids in testes and the existence of many spermatozoa in the caudal epididymis from *Lhcgr*^-/-^ mice at 4 weeks after AAV8-Lhcgr treatment (Fig. S10D, E). Together, these data support the feasibility of using our AAV vector in mice that missed puberty and suggest that adulthood might not constitute an exclusion criterion for AAV-mediated gene therapy in potential LCF candidates.

## Discussion

We herein report the first proof-of-concept study demonstrating that AAV-mediated gene therapy recovers testosterone production, restarts sexual development, rescues spermatogenesis, and restores fertility in the *Lhcgr*-deficient mouse model of LCF, suggesting that the AAV gene therapy strategy appears to be a promising treatment for LCF and thus may have value for future clinical applications.

Previous studies have shown that long-term TRT could recover serum testosterone levels and promote virilization in male pubertal *Lhcgr*^-/-^ mice ^19, 21^. When the mean testosterone levels were approximately 4- to 8-fold higher than those in the wild-type group, TRT stimulated the production of mature sperm in the treated *Lhcgr*^-/-^ mice ^19, 21^. However, exogenous testosterone beyond physical dosage can induce many serious adverse effects ^31, 32^; thus, it is not applicable in clinical practice. In this study, AAV8-mediated gene therapy in pubertal *Lhcgr*^-/-^ mice partially increased serum testosterone levels and substantially improved in sexual development. More importantly, modest testosterone restoration resulted in significant improvement in fertility. Testosterone restoration of approximately 20% in AAV8-Lhcgr-injected *Lhcgr*^-/-^ mice recovered full spermatogenesis, produced fertilization-competent spermatids, and effectively gave rise to offspring. This might be due to the recovery of intratesticular testosterone and nonsteroidal factor Insl3, which are vital for spermatogenesis ^7, 33^. Notably, these AAV8-Lhcgr gene therapy-derived mice restored fertility and produced the second generation.

Ideally, gene therapy should be offered to patients at an early disease stage ^11^. However, a large proportion of LCF patients are not diagnosed until adulthood ^28,29,30^. To our surprise, we herein found that adult mice responded to our AAV-mediated gene therapy as effectively as pubertal mice, as demonstrated by similar increments of testosterone production and restarting of sexual development. Notably, we observed many spermatids in the testis and caudal epididymis of *Lhcgr*^-/-^ mice in adult cohorts after AAV8-Lhcgr treatment. Although IVF experiments were not conducted to evaluate the fertility of *Lhcgr*^-/-^ mice treated at adult ages, we believe that these sperm would generate zygotes and fertile offspring as observed in the pubertal cohort. This finding bodes well for translation into clinical application, meeting the urgent need of patients who already have reached adulthood at the time of diagnosis and might be facing tremendous pressures related to fertility ^6^.

AAV-mediated gene therapy has been successfully applied to treat various genetic diseases in animal models ^11, 15, 34^. However, there have been relatively few studies about AAV gene transfer in testis, possibly due to safety and ethical considerations ^35, 36^. We herein showed that interstitially injected AAV8 exhibited efficient progenitor Leydig cells transduction without targeting germ cells or Sertoli cells. Similarly, Satoshi and colleagues ^36^ did not observe infected germ or Sertoli cells following interstitial injection of AAV8. Even when administered by tubular injection at high dose (1×10^13 gc/mL), AAV8 selectively transduced Sertoli cells but not germ cells. We also analyzed tail DNA from the offspring of *Lhcgr*^-/-^ mice treated with AAV8-Lhcgr and demonstrated no AAV8 integration in genomic DNA, indicating that AAV8 did not transduce germ cells or cause stable integration into the genome. Given the importance of large animal models in translational research ^37^, we further injected AAV8 particles into testes of two pubertal male cynomolgus monkeys (Macaca fascicularis). In accordance with the results in mouse experiments, AAV8 targeted progenitor Leydig cells but not germ cells or Sertoli cells in monkey testes (data not shown), reinforcing the safety and feasibility of AAV gene therapy in testis.

AAV-mediated gene delivery is an accurate etiological treatment that is ideal for genetic diseases ^13, 17^. Based on the therapeutic effects of AAV8-Lhcgr on *Lhcgr*-deficient LCF mice, it is rational to hypothesize that AAV-mediated gene therapy exhibits strong potential to exert favorable effects on other types of genetic LCF, such as 3β-hydroxysteroid dehydrogenase type 2 (3β-HSD2) deficiency ^38^, cytochrome P450 oxidoreductase (POR) deficiency ^39^, CYP17A1 deficiency ^40^, or 17β-hydroxysteroid dehydrogenase type 3 (17β-HSD3) deficiency ^41^. Although further studies are needed to thoroughly assess whether these types of LCF can be addressed by AAV8-mediated gene therapy, the present study shed new light on the novel treatment of LCF, which can potentially be expanded to more than 70 forms of genetic gonadal failure in an individualized manner ^42^.

In summary, we herein used *Lhcgr*^-/-^ male mice for initial proof-of-concept studies that established the feasibility of AAV-mediated gene therapy for genetic LCF. Treating *Lhcgr*^-/-^ mice with AAV8-Lhcgr results in robust expression of the *Lhcgr* transgene in testes, giving rise to considerable testosterone recovery and Leydig cell maturation. Notably, this therapy resulted in substantial improvement of sexual development, partial restoration of spermatogenesis, and production of fertile offspring. Furthermore, these favorable and promising effects could be reproduced in adult *Lhcgr*^-/-^ male mice treated with AAV8-Lhcgr. It is expected that these proof-of-concept data will set the stage for further studies and promote potential clinical applications of gene therapies for genetic LCF.

## Methods

### Experimental Design

Male *Lhcgr*-deficient (*Lhcgr*^-/-^) mice were injected interstitially with AAV vectors encoding *Lhcgr* at 3 weeks, 8 weeks, and 6 months of age. Male littermates expressing wild-type *Lhcgr* (*Lhcgr*^+/+^) or heterozygous for the null mutation (*Lhcgr*^+/-^) were used as controls. The therapeutic readouts included testicular *Lhcgr* expression, testosterone levels, sexual development, histological evaluation, spermatogenesis assay, and fertility analyses. Experimental groups were sized to allow for statistical analysis and the sample size for each experiment is noted in the figure legends. Mice were assigned randomly to the experimental groups, and the investigators responsible for assessing and measuring results were blinded to the intervention. Strategies used for sample collection, treatment, and information processing are included in the Results and Materials and Methods sections. Primary data are presented in data file S1.

### Animals

A breeding colony was established from *Lhcgr* heterozygous (*Lhcgr*^+/-^) C57BL/6 mice provided by Z.L. ^18^. Male mice were identified by PCR performed on DNA isolated from the tail as previously described ^18^ and were randomly assigned to experimental groups. All mice were maintained under controlled temperature (24□±□1°C) and relative humidity (50–60%) with a standard 12-hour light-and-dark cycle for the duration of the study. All animal experiments were carried out using protocols approved by the ICE for Clinical Research and Animal Trials of the First Affiliated Hospital of Sun Yat-sen University (No. 2020-003).

### Gene delivery in animal models

The full-length mouse *Lhcgr* cDNA was inserted into the pAAV-CAG vector. After sequencing-based verification, the constructed vector, called AAV8-Lhcgr, was custom packaged, purified, and titrated by Vigene Bioscience (Shandong, China). For interstitial injection of PBS or AAV vectors, we modified a previously reported method ^36^. Briefly, mice were placed under general anesthesia and the testes were exposed. Each testis was immobilized by microforceps, AAV particles or PBS were injected into the testes using a 33-gauge needle (8 μL/testis), and the incision was sutured.

### RNA extraction, cDNA synthesis, and quantitative RT-PCR

Total RNA was extracted from mouse testes using an RNeasy Kit (Qiagen, Germantown, MD, USA) according to the manufacturer’s protocol. RNA purity was determined using a NanoDrop 1000 (Thermo Fisher Scientific, Wilmington, DE, USA). Reverse transcription was performed using a NovoScript® 1st Strand cDNA Synthesis Kit (Novoprotein, Shanghai, China). Quantitative RT-PCR was performed using LightCycle®480 SYBR Green I Master (Roche, Indianapolis, IN, USA), on a Light Cycler 480 Detection System (Roche). The primers used for quantitative RT-PCR are listed in Table S2. To validate the primers, a melting curve was generated to confirm a single peak and rule out the possibility of non-specific product or primer dimer formation. β-actin was amplified as a control, and the target gene expression levels were calculated using the ΔCt method and expressed relative to β-actin.

### Immunofluorescence staining

Immunofluorescence staining was conducted as previously reported by our group ^43^. The harvested testes were fixed with 4% PFA for 2 hours at 4□. The tissues were then dehydrated with 30% sucrose solution at 4°C for 24 h. Afterwards, the tissues were soaked in Tissue-Tek O.C.T. Compound (Sakura Finetek, Torrance, CA, USA), frozen, and cryosectioned at 10 μm thickness using a frozen slicer (Leica CM1950). For intracellular protein detection, sections were permeabilized with 0.3% Triton X-100 (Sigma-Aldrich) for 30 min. The non-specific binding of antibodies was blocked with 3% BSA (Sigma-Aldrich) for 45 min at room temperature, and the slices were incubated overnight with primary antibodies at 4°C. The sections were thereafter washed with PBS for 3 times and incubated with secondary antibodies at room temperature for 45 min in the dark, and then for 5 min with DAPI. Finally, the specific fluorescence was visualized and photographed using an LSM800 confocal microscope (Zeiss, Jena, Germany) or a Leica DMi8 microscope (Leica, Wetzlar, Germany). The primary and secondary antibodies are listed in Table S3.

### Histological analysis

The testis and epididymis were collected, fixed overnight in Bouin’s solution (Sigma-Aldrich), dehydrated in 75% ethanol, embedded in paraffin, and sectioned at 4 μm. The sections were deparaffinized with xylene, hydrated with graded ethanol, and stained with hematoxylin and eosin for histological analysis using an AxioScan.Z1 (Zeiss) or a Leica DMi8 microscope (Leica).

### Sex hormone assays

Sex hormone concentrations were assayed as previously reported by our group ^44^. Serum and testes samples were collected at the indicated timepoints and stored at -80°C until analysis. Testosterone levels were measured using a chemiluminescent immunoassay (CLIA) (KingMed Diagnostics Group Co., Ltd. Guangzhou, China). The coefficient of variation (CV) of CLIA is 2-5.1% for intra-assay precision and 2.6-5.2% for inter-assay precision. The minimum detectable dose of testosterone is 0.01 ng/mL.

### Computer-aided semen analysis

Semen samples were analyzed as previously reported ^27^. One cauda epididymis was removed from each mouse, incised with micro scissors, and incubated in 0.5 mL buffer containing 0.5% BSA (Sigma) for 15 min at 37°C to allow for sperm release. The tissue was removed, and sperm samples were diluted and analyzed using a Hamilton Thorne’s Ceros II system. At least six fields were assessed for each sample, and the sperm concentration and percentages of motile and progressively motile spermatozoa were determined.

### RNA-Seq analysis

Total mRNA was isolated from the testes of *Lhcgr*^+/+^ mice, *Lhcgr*^+/-^ mice, and *Lhcgr*^-/-^ mice injected with PBS or AAV8-Lhcgr. RNA sequencing libraries were constructed using an Illumina mRNA-seq Prep Kit (Illumina, San Diego, CA, USA) as recommended by the manufacturer. The fragmented and randomly primed 150-bp paired-end libraries were sequenced using an Illumina HiSeq X Ten. The sequencing data were processed using Consensus Assessment of Sequence and Variation (CASAVA, version 1.8.2; Illumina) under the default settings. The transcripts per million (TPM) values were used to evaluate the expression levels of genes. Pearson correlation coefficient (PCC) values and heat maps of the global gene expression profiles were generated to measure the similarities of the different samples. Differentially expressed genes (DEGs) between PBS-treated and AAV-Lhcgr-treated *Lhcgr*^-/-^ testes were analyzed, and the relevant biological processes were identified by Gene Ontology (GO) analysis. Finally, the RNA-Seq data were analyzed under R version 4.0.2 to categorize the defined genes that characterized spermatogonia, spermatocytes, round spermatocytes, and elongating spermatids ^25, 26^.

### In vitro fertilization (IVF) and mouse embryo transfer

IVF was conducted as previously described ^45^. Sperm were collected from the caudal epididymis of *Lhcgr*^-/-^ mice (7 weeks old) at 4 weeks after AAV8-*Lhcgr* injection, and then incubated in a humidified atmosphere of 5% CO2 in air at 37°C for 2 hours in a 200 μL drop of TYH medium covered with paraffin oil (Sigma-Aldrich) for sperm capacitation. Cumulus-oocyte complexes were collected from the oviduct of wild-type female C57BL/6 mice at 16 h after ovulatory human chorionic gonadotropin (hCG, Sigma-Aldrich) injection and placed in a 50 µL drop of TYH medium that was then covered with mineral oil (Sigma-Aldrich). IVF was performed by the addition of in vitro-capacitated sperm (approximately 10,000 sperm/50 µL) into the medium containing the oocytes. After 4 h, the oocytes were collected and transferred to the developing medium, KSOM (Sigma-Aldrich). All zygotes were then washed and transferred into KSOM (Sigma-Aldrich), and cultured in a humidified atmosphere of 5% CO2 in air at 37°C. The numbers of 2-cell embryos were determined at 21 hours after the insemination. The two-cell embryo formation rate was calculated by dividing the number of 2-cell embryos by the number of zygotes examined. For embryo transfer, we mated 8- to 12-week-old females with vasectomized males and determine whether they are pseudo-pregnant by the presence of a vaginal plug. 2-cell embryos were surgically transferred into the uterine horn of each pseudo-pregnant female.

### PCR analysis of AAV8-Lhcgr integration in the offspring

To test whether AAV8 integrated into the genome of offspring, we applied a previously reported method ^36^. Total DNA was extracted from the tail using a TIANamp Genomic DNA kit (TIANGEN, Beijing, China) according to the manufacturer’s instructions. Genome integrations were quantified by PCR using specific primers for the CAG promoter and the inserted *Lhcgr*. PCR was performed using a Bio-Rad T100 following the manufacturer’s protocol. The primers used for PCR are listed in Table S2.

### Statistical Analysis

All data were analyzed using the SPSS 20.0 software (IBM SPSS Statistics, Armonk, NY, USA). Statistical differences between samples were assessed with Student’s t-tests, one-way analysis of variance (ANOVA), or nonparametric tests (Mann-Whitney U tests and Kruskal-Wallis test). Differences were considered significant when P < 0.05 (*P < 0.05, **P < 0.01 and ***P < 0.001).

## Supporting information

Primary Data

## Acknowledgments

The authors thank the Core Facilities of Medical Science for support in imaging experiments and are grateful to Yan Guo for her constructive suggestions, Rongda Deng for generously providing advice on testosterone detection, Zexin Guo for his assistance in semen analyses, and Yuan Qiu for her support of animal experiments.

## Author contributions

K.X., F.W. and X.L. contributed equally to this work. K.X. carried out the experiments, assisted with the experimental design and wrote the manuscript. F.W. carried out the experiments and data analysis. X.L. assisted with the design of the viral vector and assisted with the experimental design. P.L. performed animal experiments. H.C. assisted with IVF and data analysis. Y.M. and W.H. provided technical help in RNA-seq analysis. W.O., Y.L. and X.F. assisted with animal experiments. Z.L. provided *Lhcgr*^-/-^ mice and assisted with manuscript revision. X.T., Q.K. and F.F.M. assisted with the experimental design and revised the manuscript. A.P.X. and C.D. conceived the project, supervised all experiments, and wrote and revised the manuscript. All authors fulfil the criteria for authorship.

## Data and materials availability

Reagents and mouse models described here are accessible through a material transfer agreement. All data associated with this study are presented in the paper or the Supplementary Materials.

## References

1. Salonia A, et al. Paediatric and adult-onset male hypogonadism. Nat Rev Dis Primers 5, 38 (2019).

2. Zirkin BR, Papadopoulos V. Leydig cells: formation, function, and regulation. Biol Reprod 99, 101–111 (2018).

3. Mendonca BB, Costa EM, Belgorosky A, Rivarola MA, Domenice S. 46,XY DSD due to impaired androgen production. Best Pract Res Clin Endocrinol Metab 24, 243–262 (2010).

4. Teerds KJ, Huhtaniemi IT. Morphological and functional maturation of Leydig cells: from rodent models to primates. Hum Reprod Update 21, 310–328 (2015).

5. Bhasin S, et al. Testosterone Therapy in Men With Hypogonadism: An Endocrine Society Clinical Practice Guideline. J Clin Endocrinol Metab 103, 1715–1744 (2018).

6. Guercio G, Costanzo M, Grinspon RP, Rey RA. Fertility Issues in Disorders of Sex Development. Endocrinol Metab Clin North Am 44, 867–881 (2015).

7. Kathrins M, Niederberger C. Diagnosis and treatment of infertility-related male hormonal dysfunction. Nat Rev Urol 13, 309–323 (2016).

8. Samaranch L, et al. Adeno-associated viral vector serotype 9-based gene therapy for Niemann-Pick disease type A. Sci Transl Med 11, eaat3738 (2019).

9. Song Y, et al. Non-immunogenic utrophin gene therapy for the treatment of muscular dystrophy animal models. Nat Med 25, 1505–1511 (2019).

10. Cehajic-Kapetanovic J, et al. Initial results from a first-in-human gene therapy trial on X-linked retinitis pigmentosa caused by mutations in RPGR. Nat Med 26, 354–359 (2020).

11. Manso AM, et al. Systemic AAV9. LAMP2B injection reverses metabolic and physiologic multiorgan dysfunction in a murine model of Danon disease. Sci Transl Med 12, eaax1744 (2020).

12. Pasi KJ, et al. Multiyear Follow-up of AAV5-hFVIII-SQ Gene Therapy for Hemophilia A. N Engl J Med 382, 29–40 (2020).

13. Wang D, Tai PWL, Gao G. Adeno-associated virus vector as a platform for gene therapy delivery. Nat Rev Drug Discov 18, 358–378 (2019).

14. Mendell JR, et al. Assessment of Systemic Delivery of rAAVrh74.MHCK7.micro-dystrophin in Children With Duchenne Muscular Dystrophy: A Nonrandomized Controlled Trial. JAMA Neurol 77, 1122–1131 (2020).

15. Wang S, et al. AAV Gene Therapy Prevents and Reverses Heart Failure in a Murine Knockout Model of Barth Syndrome. Circ Res 126, 1024–1039 (2020).

16. Pan B, et al. Gene therapy restores auditory and vestibular function in a mouse model of Usher syndrome type 1c. Nat Biotechnol 35, 264–272 (2017).

17. Li CW, Samulski RJ. Engineering adeno-associated virus vectors for gene therapy. Nat Rev Genet 21, 255–272 (2020).

18. Lei ZM, et al. Targeted disruption of luteinizing hormone/human chorionic gonadotropin receptor gene. Mol Endocrinol 15, 184–200 (2001).

19. Pakarainen T, Zhang FP, Makela S, Poutanen M, Huhtaniemi I. Testosterone replacement therapy induces spermatogenesis and partially restores fertility in luteinizing hormone receptor knockout mice. Endocrinology 146, 596–606 (2005).

20. Rahman NA, Rao CV. Recent progress in luteinizing hormone/human chorionic gonadotrophin hormone research. Mol Hum Reprod 15, 703–711 (2009).

21. Lei ZM, Mishra S, Ponnuru P, Li X, Yang ZW, Rao Ch V. Testicular phenotype in luteinizing hormone receptor knockout animals and the effect of testosterone replacement therapy. Biol Reprod 71, 1605–1613 (2004).

22. Kossack N, Troppmann B, Richter-Unruh A, Kleinau G, Gromoll J. Aberrant transcription of the LHCGR gene caused by a mutation in exon 6A leads to Leydig cell hypoplasia type II. Mol Cell Endocrinol 366, 59–67 (2013).

23. Zhang FP, Pakarainen T, Zhu F, Poutanen M, Huhtaniemi I. Molecular characterization of postnatal development of testicular steroidogenesis in luteinizing hormone receptor knockout mice. Endocrinology 145, 1453–1463 (2004).

24. Welsh M, et al. Identification in rats of a programming window for reproductive tract masculinization, disruption of which leads to hypospadias and cryptorchidism. J Clin Invest 118, 1479–1490 (2008).

25. Ernst C, Eling N, Martinez-Jimenez CP, Marioni JC, Odom DT. Staged developmental mapping and X chromosome transcriptional dynamics during mouse spermatogenesis. Nat Commun 10, 1251 (2019).

26. Green CD, et al. A Comprehensive Roadmap of Murine Spermatogenesis Defined by Single-Cell RNA-Seq. Dev Cell 46, 651–667 e610 (2018).

27. Matzuk MM, et al. Small-molecule inhibition of BRDT for male contraception. Cell 150, 673–684 (2012).

28. Martens JW, et al. A homozygous mutation in the luteinizing hormone receptor causes partial Leydig cell hypoplasia: correlation between receptor activity and phenotype. Mol Endocrinol 12, 775–784 (1998).

29. Troppmann B, Kleinau G, Krause G, Gromoll J. Structural and functional plasticity of the luteinizing hormone/choriogonadotrophin receptor. Hum Reprod Update 19, 583–602 (2013).

30. Bakircioglu ME, Tulay P, Findikli N, Erzik B, Gultomruk M, Bahceci M. Successful testicular sperm recovery and IVF treatment in a man with Leydig cell hypoplasia. J Assist Reprod Genet 31, 817–821 (2014).

31. Gagliano-Juca T, Basaria S. Testosterone replacement therapy and cardiovascular risk. Nat Rev Cardiol 16, 555–574 (2019).

32. Tsametis CP, Isidori AM. Testosterone replacement therapy: For whom, when and how? Metabolism 86, 69–78 (2018).

33. Sagata D, et al. The Insulin-Like Factor 3 (INSL3)-Receptor (RXFP2) Network Functions as a Germ Cell Survival/Anti-Apoptotic Factor in Boar Testes. Endocrinology 156, 1523–1539 (2015).

34. Weber ND, et al. Gene therapy for progressive familial intrahepatic cholestasis type 3 in a clinically relevant mouse model. Nat Commun 10, 5694 (2019).

35. Rajasekaran S, et al. Infectivity of adeno-associated virus serotypes in mouse testis. BMC Biotechnol 18, 70 (2018).

36. Watanabe S, Kanatsu-Shinohara M, Ogonuki N, Matoba S, Ogura A, Shinohara T. In Vivo Genetic Manipulation of Spermatogonial Stem Cells and Their Microenvironment by Adeno-Associated Viruses. Stem Cell Reports 10, 1551–1564 (2018).

37. Knechtle SJ, Shaw JM, Hering BJ, Kraemer K, Madsen JC. Translational impact of NIH-funded nonhuman primate research in transplantation. Sci Transl Med 11, eaau0143 (2019).

38. Burckhardt MA, et al. Human 3beta-hydroxysteroid dehydrogenase deficiency seems to affect fertility but may not harbor a tumor risk: lesson from an experiment of nature. Eur J Endocrinol 173, K1–K12 (2015).

39. Miller WL. P450 oxidoreductase deficiency: a disorder of steroidogenesis with multiple clinical manifestations. Sci Signal 5, pt11 (2012).

40. Marsh CA, Auchus RJ. Fertility in patients with genetic deficiencies of cytochrome P450c17 (CYP17A1): combined 17-hydroxylase/17,20-lyase deficiency and isolated 17,20-lyase deficiency. Fertil Steril 101, 317–322 (2014).

41. Yang Z, et al. 17beta-Hydroxysteroid dehydrogenase 3 deficiency: Three case reports and a systematic review. J Steroid Biochem Mol Biol 174, 141–145 (2017).

42. Wisniewski AB, et al. Management of 46,XY Differences/Disorders of Sex Development (DSD) Throughout Life. Endocr Rev 40, 1547–1572 (2019).

43. Xia K, et al. Endosialin defines human stem Leydig cells with regenerative potential. Hum Reprod 35, 2197–2212 (2020).

44. Xia K, et al. Restorative functions of Autologous Stem Leydig Cell transplantation in a Testosterone-deficient non-human primate model. Theranostics 10, 8705–8720 (2020).

45. Umehara T, Tsujita N, Zhu ZD, Ikedo M, Shimada M. A simple sperm-sexing method that activates TLR7/8 on X sperm for the efficient production of sexed mouse or cattle embryos. Nature Protocols 15, 2645–2667 (2020).

