## Supplemental Material for "AAV-mediated gene therapy produces fertile offspring in the *Lhcgr*-deficient mouse model of Leydig cell failure"

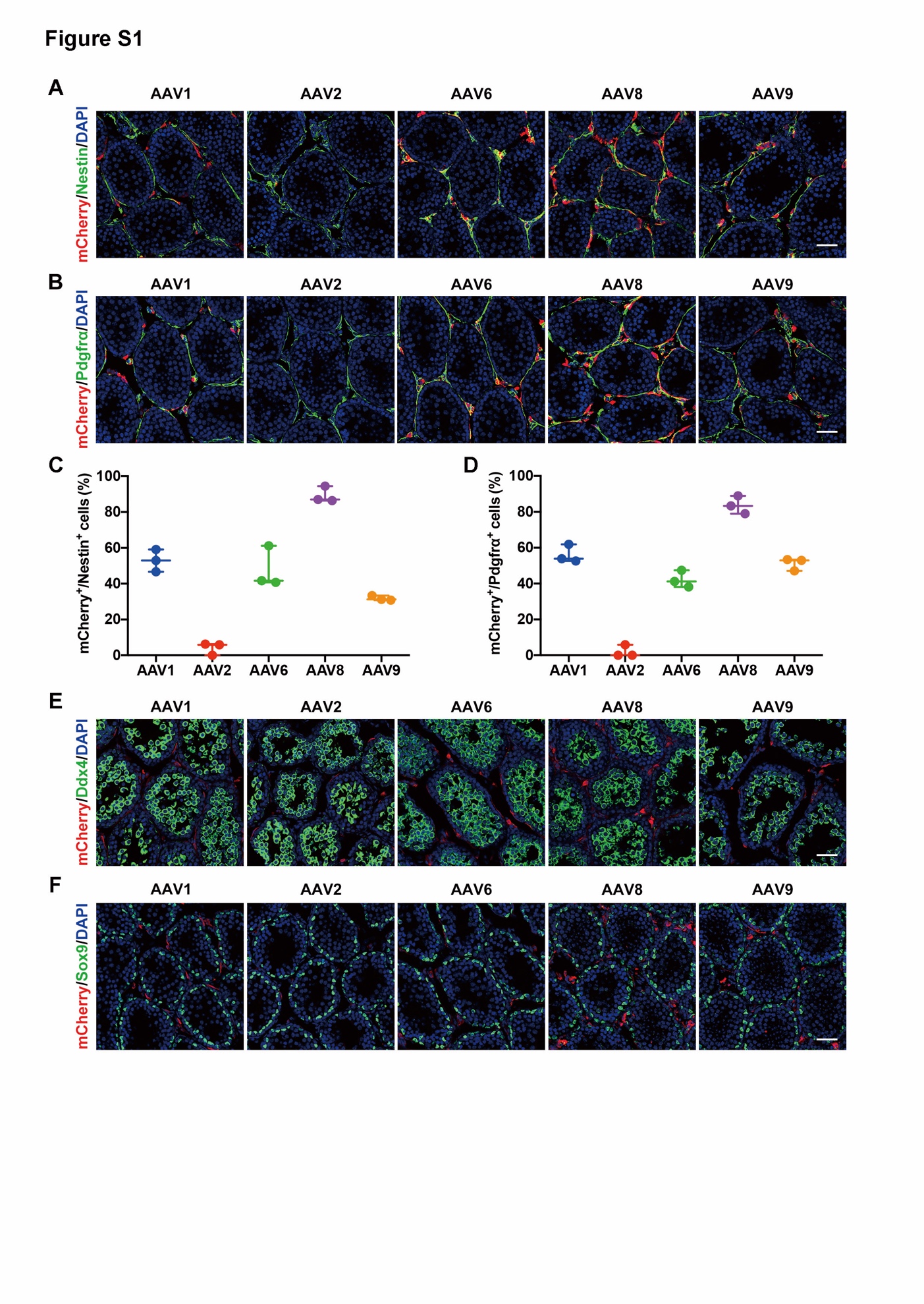

**Fig. S1. Testicularly injected AAV8 targets progenitor Leydig cells**

(A,B) Representative confocal images of the testicular sections of *Lhcgr*^-/-^ mice injected with AAV-CAG-mCherry (8×10^10 gc/testis, n=3) of the indicated capsid serotypes. The testis tissues were collected and immunostained with progenitor Leydig cells markers (Nestin, Pdgfrα) at 7 days after AAV injection. Scale bars: 50 μm. (C,D) Viral transduction rates were determined from the number of mCherry^+^ cells divided by the number of Nestin^+^ or Pdgfrα^+^ progenitor Leydig cells. Data are represented by plots, and whiskers are minimum to maximum values. (E,F) Immunostaining of AAV-mCherry-injected testes for the germ cell marker, Ddx4, or the Sertoli cell marker, Sox9, at 7 days after injection. Nuclei were counterstained with DAPI. Scale bar: 50 μm.

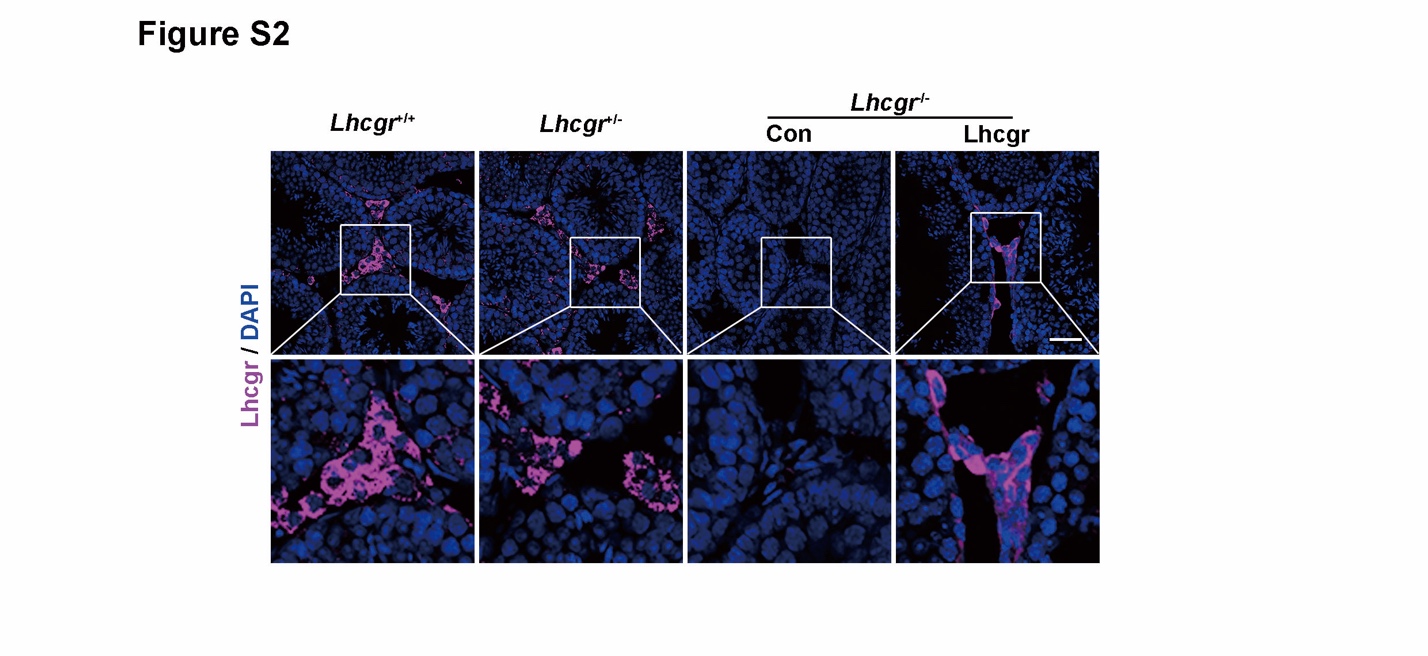

**Fig. S2. Lhcgr expression in the testes of recipient mice in pubertal cohort**

Representative images of *Lhcgr* expression in the testicular interstitium at 4 weeks after treatment in *Lhcgr*^+/+^ mice (n=4), *Lhcgr*^+/-^ mice (n=4), and *Lhcgr*^-/-^ mice injected with PBS (n=4) or AAV8-Lhcgr (8×10^10 gc/testis, n=4). Nuclei were counterstained with DAPI. Scale bar: 100 μm.

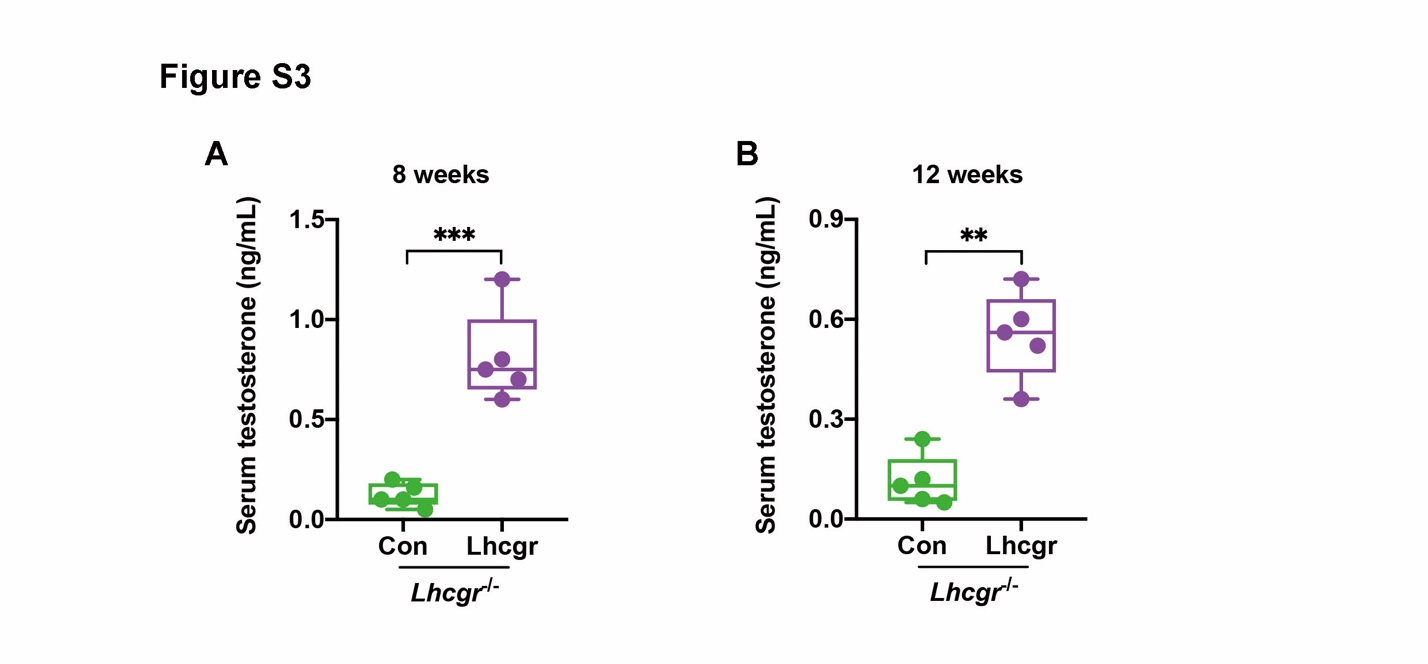

**Fig. S3. Serum testosterone levels after gene therapy**

(A,B) The concentrations of serum testosterone were analyzed 8 (A) and 12 weeks (B) after treatment in *Lhcgr*^-/-^ mice injected with PBS (n=5) or AAV8-Lhcgr (8×10^10 gc/testis, n=5). Data are represented by box plots, and whiskers are minimum to maximum values. Significance was determined by Student’s t-test. ** P < 0.01, *** P < 0.001.

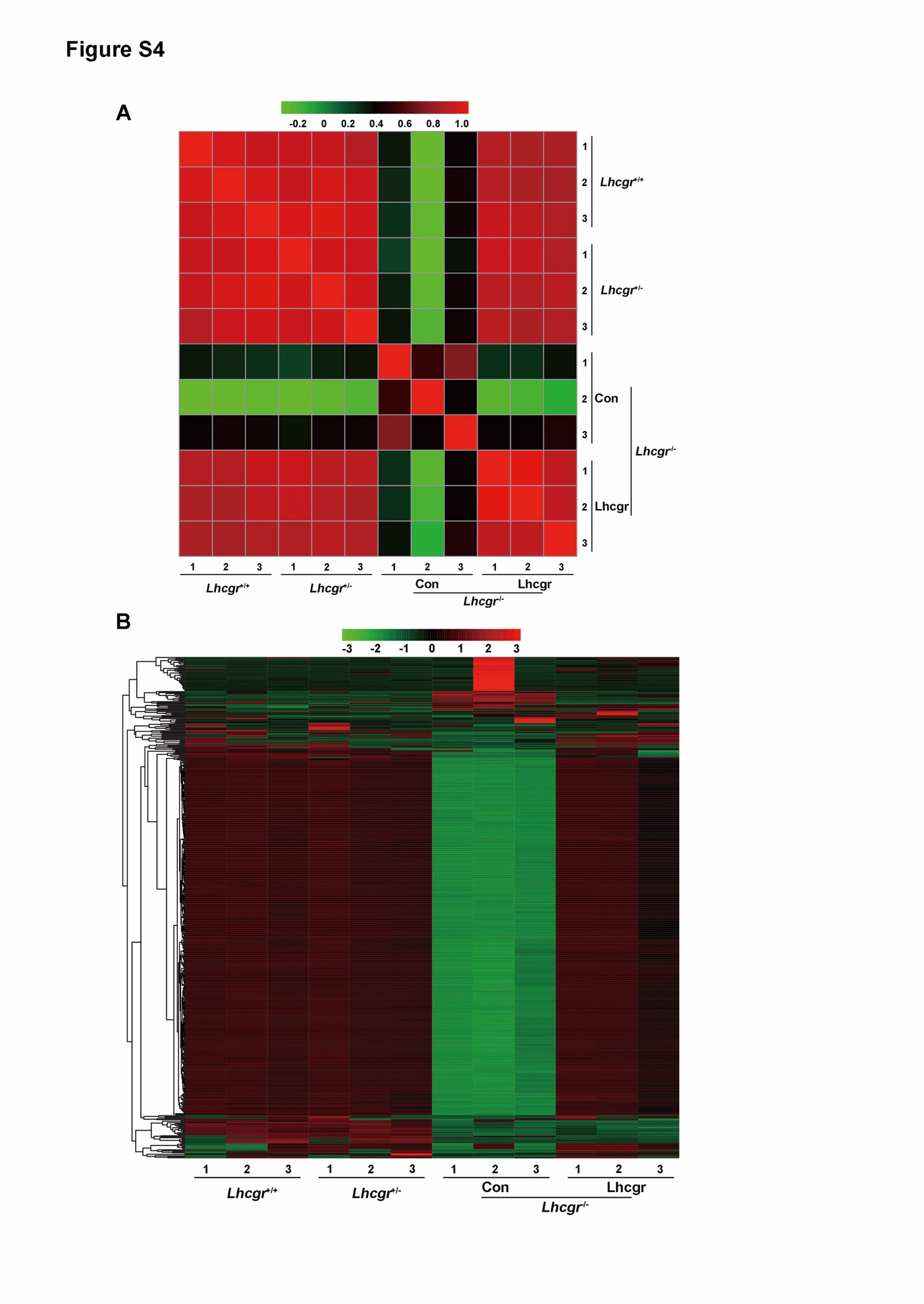

**Fig. S4. Pearson correlation coefficient and heatmap of** **the testicular transcriptome profiles**

(A) Pearson correlation coefficient (PCC) values were calculated to identify the correlation among *Lhcgr*^+/+^ mice (n=3), *Lhcgr*^+/-^ mice (n=3), and *Lhcgr*^-/-^ mice injected with PBS (n=3) or AAV8-Lhcgr (8×10^10 gc/testis, n=3). Each symbol represents an individual sample. (B) Heat map of top 500 gene expression profiles in the testes of *Lhcgr*^+/+^ mice (n=3), *Lhcgr*^+/-^ mice (n=3), and *Lhcgr*^-/-^ mice injected with PBS (n=3) or AAV8-Lhcgr (8×10^10 gc/testis, n=3). The color code indicates z-score-normalized expression values.

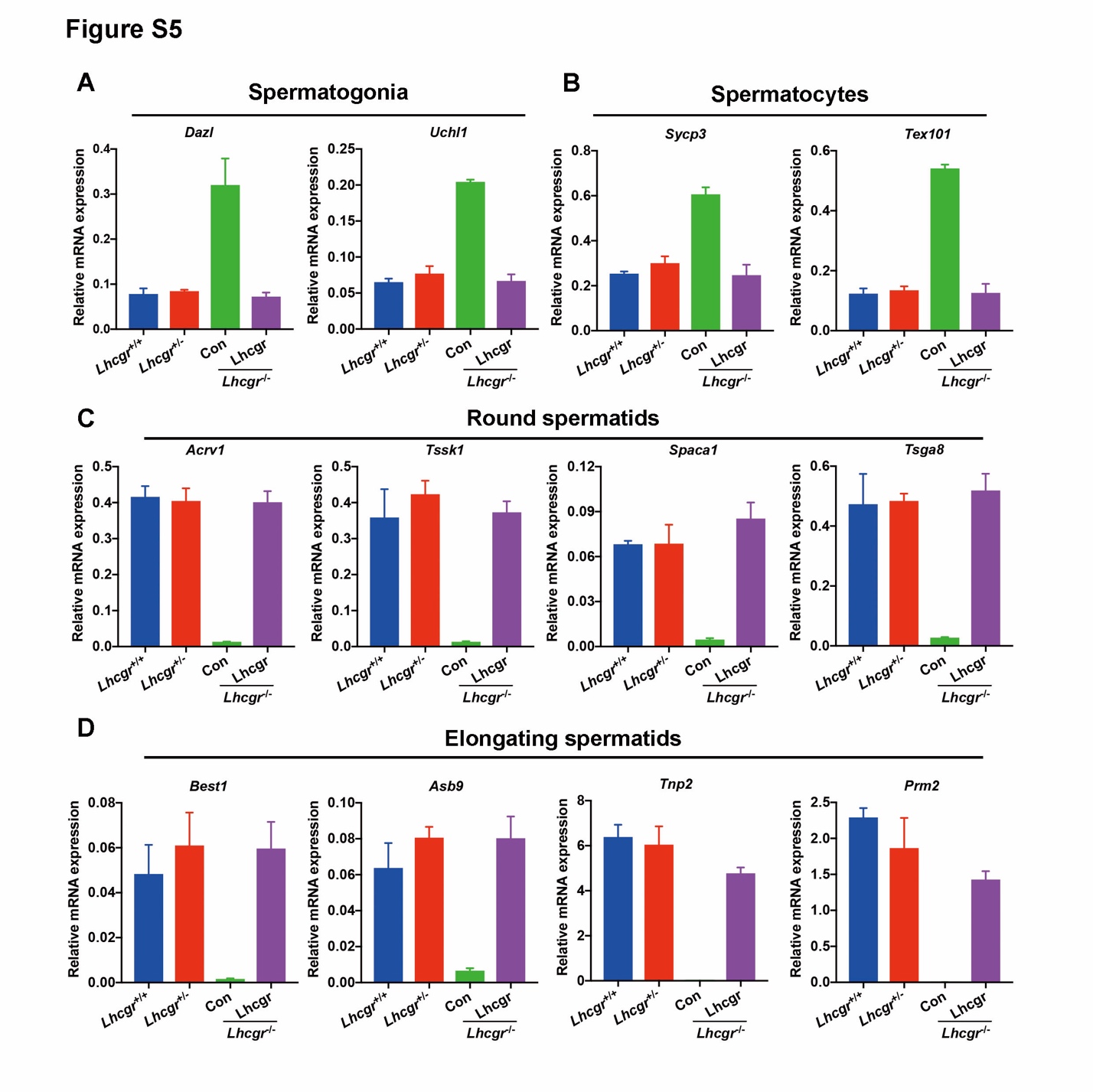

**Fig. S5. mRNA expression of specific germ cell marker genes, as analyzed by quantitative RT-PCR**

(A-D) Quantitative RT-PCR analysis was performed in testes samples collected from *Lhcgr*^+/+^ mice (n=3), *Lhcgr*^+/-^ mice (n=3), and *Lhcgr*^-/-^ mice injected with PBS (n=3) or AAV8-Lhcgr (8×10^10 gc/testis, n=3) at 4 weeks after treatment. The expression levels of marker genes for spermatogonia (Dazl, Uchl1), spermatocytes (Tex101, Sycp3), round spermatids (Acrv1, tssk1, Spaca1, and Tsga8), and elongating spermatids (Best1, Prm2, Tnp2, and Asb9) were detected from each group. Data are expressed as mean ± sem.

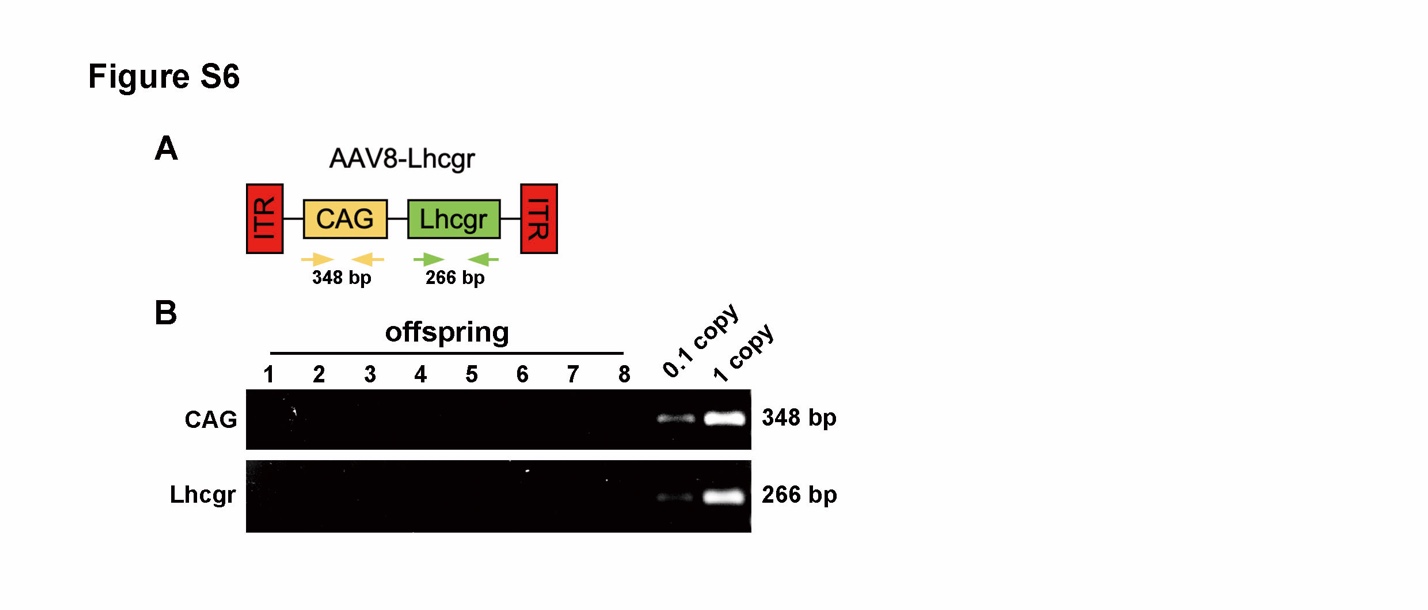

**Fig. S6.** **Analysis of AAV8 integration in offspring**

(A,B) PCR analysis of AAV8-Lhcgr integration in the genomes of offspring. CAG promoter- and *Lhcgr*-specific primers were used. Samples were collected from offspring produced after interstitial injection with AAV8-Lhcgr. As a control, tail DNA from *Lhcgr*^+/-^ mice was spiked with viral particles representing 0.1 and 1 copies of the viral genome.

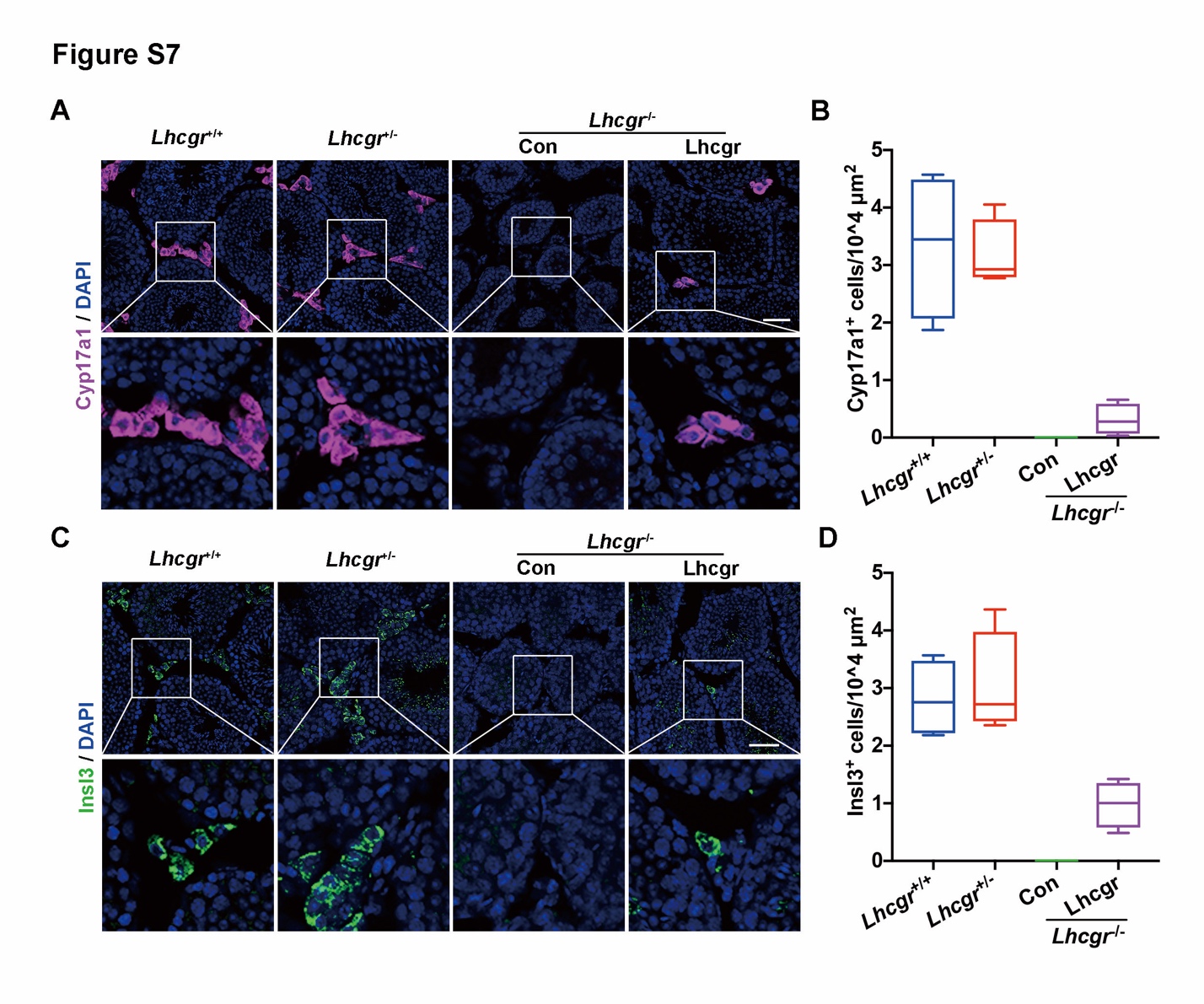

**Fig. S7. Testicular injection of AAV8-Lhcgr rescues Leydig cell function in 8-week-old adult *Lhcgr*^-/-^ mice**

(A) The Leydig cells marker, Cyp17a1, was evaluated in the testicular interstitium by immunofluorescence assay at 4 weeks after treatment in *Lhcgr*^+/+^ mice (n=4), *Lhcgr*^+/-^ mice (n=4), and *Lhcgr*^-/-^ mice injected with PBS (n=4) or AAV8-Lhcgr (8×10^10 gc/testis, n=4). Scale bar: 100 μm. (B) Cyp17a1^+^ cells (n=4) were quantified in the different groups. (C) Immunostaining with anti-Insl3 antibody was used to detect mature Leydig cells at 4 weeks after treatment in *Lhcgr*^+/+^ mice (n=4), *Lhcgr*^+/-^ mice (n=4), and *Lhcgr*^-/-^ mice injected with PBS (n=4) or AAV8-Lhcgr (8×10^10 gc/testis, n=4). Scale bar: 100 μm. (D) Insl3^+^ cells (n=4) were quantified in the different groups. Data are represented by box plots, and whiskers are minimum to maximum values.

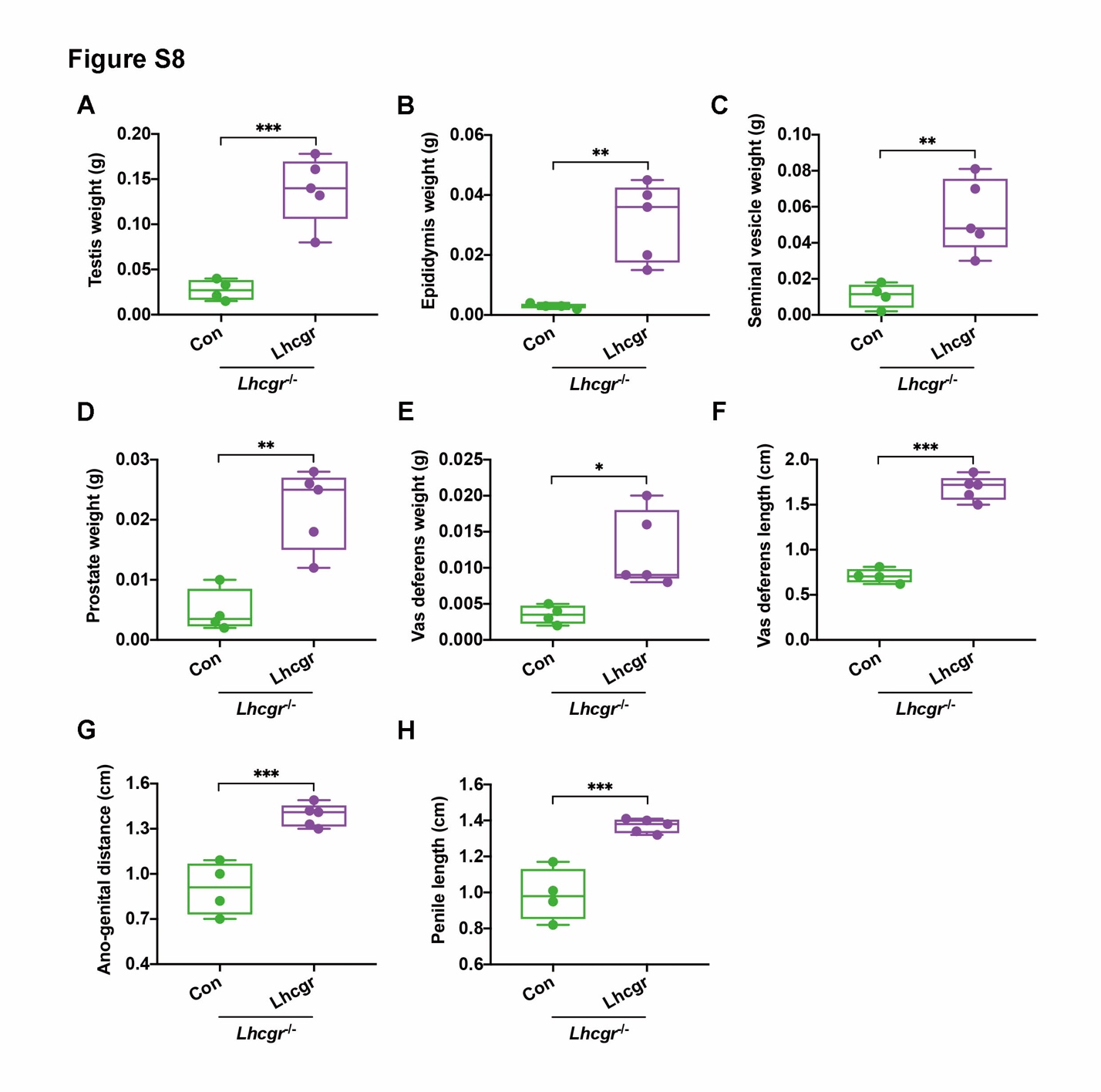

**Fig. S8. AAV8-Lhcgr restarts sexual development** **in 8-week-old adult *Lhcgr*^-/-^ mice**

(A-H) The testis weight (A), epididymis weight (B), seminal vesicle weight (C), prostate weight (D), vas deferens weight (E), vas deferens length (F), ano-genital distance (G), and penile length (H) of *Lhcgr*^-/-^ mice injected with PBS (n=4) or AAV8-Lhcgr (8×10^10 gc/testis, n=5) at 4 weeks after treatment. Data are represented by box plots, and whiskers are minimum to maximum values. Significance was determined by Student’s t-test. * P < 0.05, ** P < 0.01, *** P < 0.001.

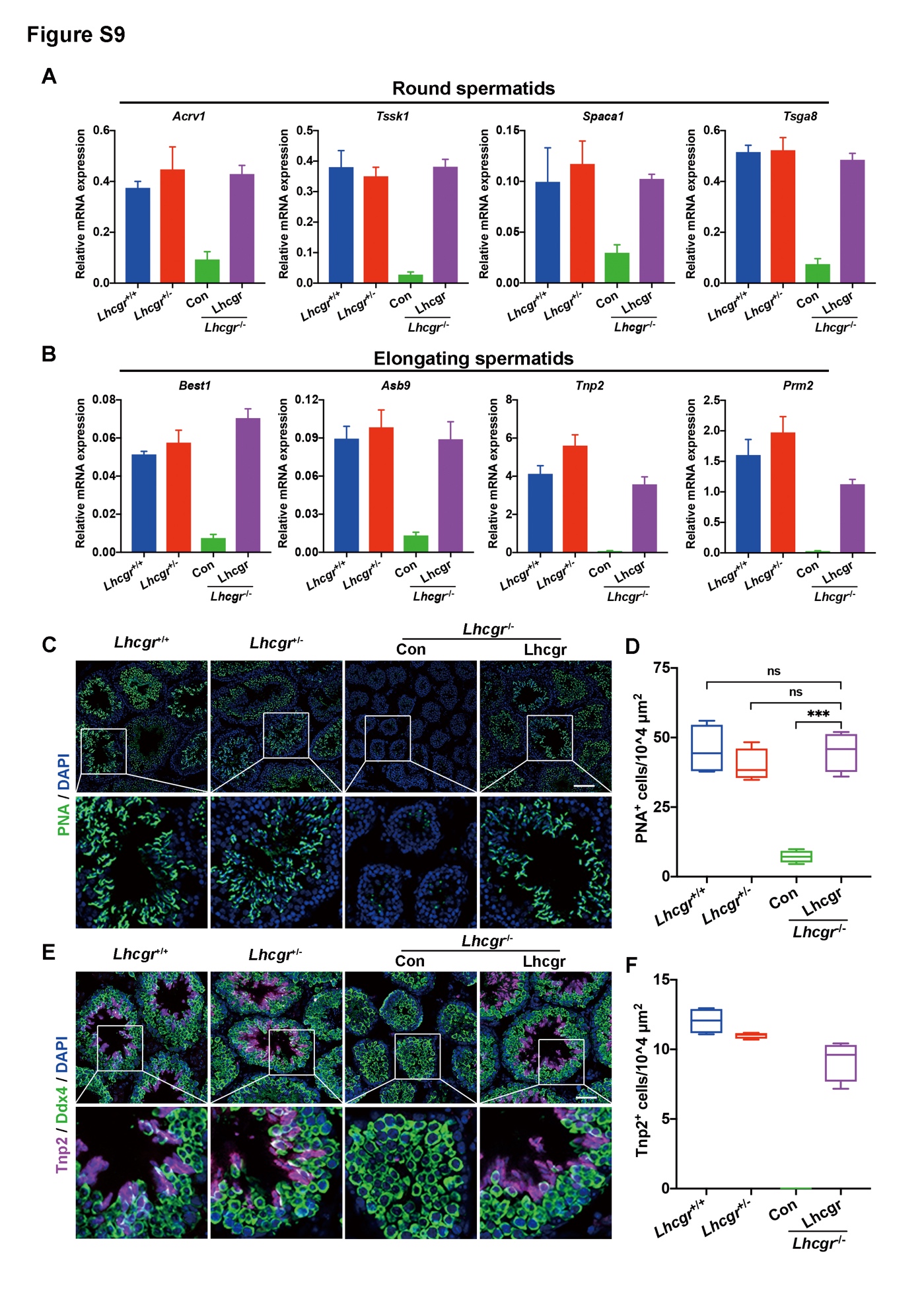

**Fig. S9. AAV8-Lhcgr promotes the** **formation of elongating spermatids in 8-week-old adult *Lhcgr*^-/-^ mice**

(A,B) Quantitative RT-PCR was used to analyze marker expression in testes samples collected from *Lhcgr*^+/+^ mice (n=3), *Lhcgr*^+/-^ mice (n=3), and *Lhcgr*^-/-^ mice injected with PBS (n=3) or AAV8-Lhcgr (8×10^10 gc/testis, n=3) at 4 weeks after treatment. The expression levels of marker genes for round spermatids (Acrv1, tssk1, Spaca1, and Tsga8) and elongating spermatids (Best1, Prm2, Tnp2, and Asb9) were detected for each group. Data are expressed as mean ± sem. (C-F) Representative images of testis sections from *Lhcgr*^+/+^ mice (n=4), *Lhcgr*^+/-^ mice (n=4), and *Lhcgr*^-/-^ mice injected with PBS (n=4) or AAV8-Lhcgr (8×10^10 gc/testis, n=4). Sections were immunostained for PNA (C), Ddx4 (E), and Tnp2 (E), and counterstained with DAPI. Quantitative analysis showing the percentages of PNA^+^ (D) and Tnp2^+^ (F) germ cells in the seminiferous tubules of the testes. Scale bars: 50 μm. Data are represented by box plots, and whiskers are minimum to maximum values. Significance was determined by one-way ANOVA. *** P < 0.001, ns = not significant.

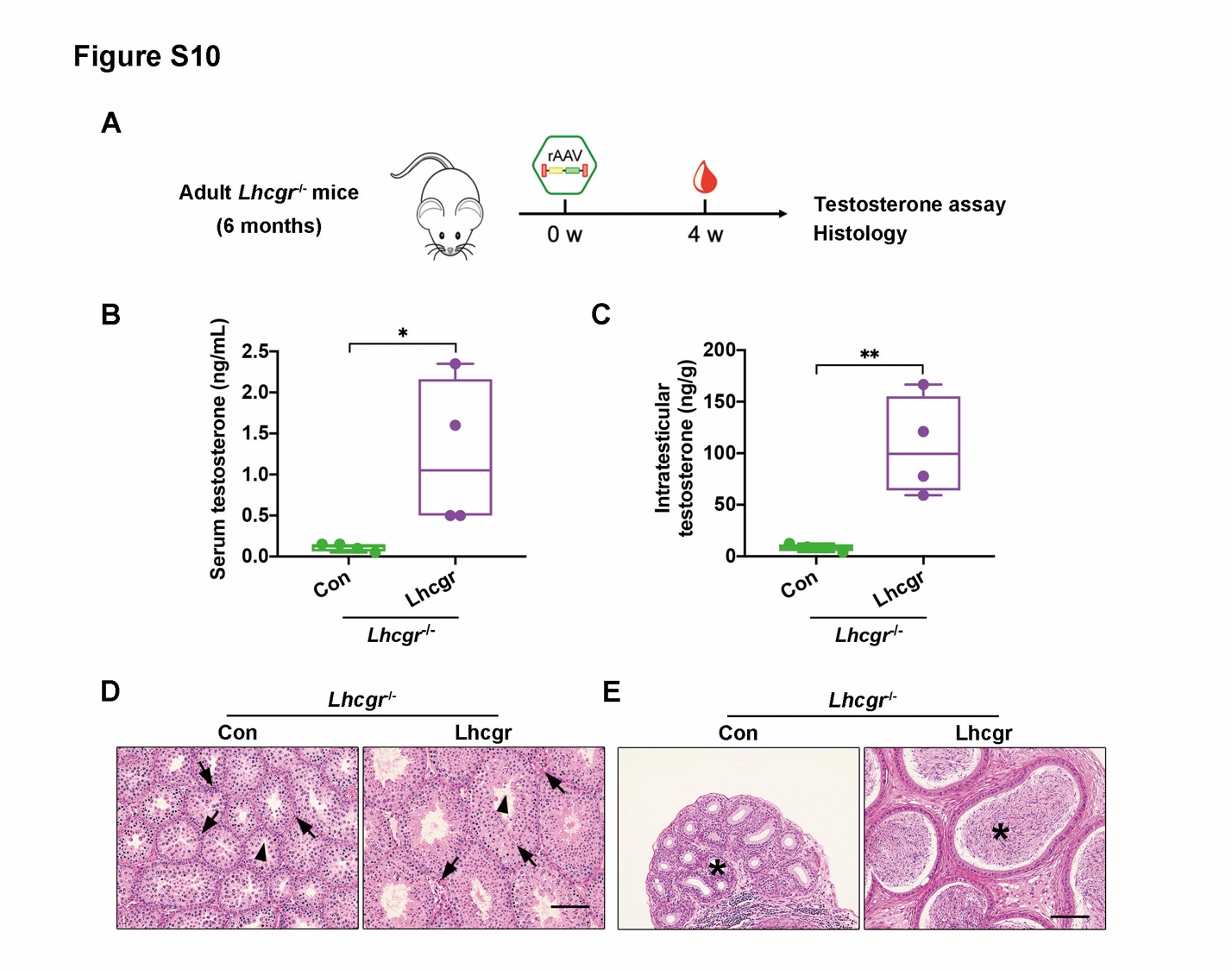

**Fig. S10. AAV8-Lhcgr recovers testosterone production and rescues spermatogenesis in 6-month-old adult *Lhcgr*^-/-^ mice**

(A) Experimental overview for the in vivo studies. (B,C) The concentrations of serum (C) and intratesticular (D) testosterone were analyzed 4 weeks after treatment in 6 months old *Lhcgr*^-/-^ mice injected with PBS (n=4) or AAV8-Lhcgr (8×10^10gc/testis, n=4). (D,E) Representative light micrographs of testes sections (D) and cauda epididymis (E) from *Lhcgr*^-/-^ mice injected with PBS (n=4) or AAV8-Lhcgr (8×10^10 gc/testis, n=4). Samples were taken 4 weeks after treatment. Arrows indicates Leydig cells and arrowheads indicates full spermatogenesis in testis (D). Stars indicates spermatozoa in the cauda epididymis (E). Scale bars: 100 μm. Data are represented by box plots, and whiskers are minimum to maximum values. Significance was determined by Student’s t-test. * P < 0.05, ** P < 0.01.

**Table S1. Development of embryos generated via IVF with sperm from AAV-Lhcgr-treated *Lhcgr*^-/-^ mice in pubertal cohort.**

| ***Lhcgr*-/- male mice** | **No. of oocytes** | **No. of 2-cell (%)** | **No. of embryos transferred** | **No. of pseudo-pregnant mice** | **No. of pups (%)** |
| --- | --- | --- | --- | --- | --- |
| M03 | 139 | 59 (42.4%) | 40 | 2 | 20 (50%) |
|  | 79 | 31 (39.2%) | 31 | 2 | 3 (9.6%) |
| M18 | 144 | 18 (12.5%) | 18 | 1 | 10 (55.6%) |
|  | 361 | 70 (19.4%) | 60 | 3 | 25 (41.7%) |

**Table S2. Primers used to amplify the transcripts in PCR analysis.**

| **Primers for Quantitative RT-PCR** | | |
| --- | --- | --- |
| **Gene** | **Forward Primer** | **Reverse Primer** |
| *Lhcgr* | CACTCTCCAGAGTTGTCAGGG | GAGGTTTGTAAAAGCACCGGG |
| *Dazl* | CTTCATCAGCAACCACAA | TTCATCCATCCTAACATCAAT |
| *Uchl1* | TGGAATTTGAGGATGGAT | AACACTTGGCTCTATCTT |
| *Sycp3* | CGCTGAGCAAACATCTAAAGA | CAACCAAAGGTGGCTTCC |
| *Tex101* | TACCTTTAACTGGACTTCA | CCATCTGCTTTAATCAACA |
| *Acrv1* | CAGGTGAACAGGTGTCTA | CAGATGTGCTTGGAAGTG |
| *Tssk1* | CAAGGACTTCAACATCAA | GGTCTTGCTTAATATCAGT |
| *Spaca1* | ATTCACCGTCTATACAAC | CAGATAATGACTCCTATGG |
| *Tsga8* | GTGAAGCCTATAATGCCAAT | ACCCTTTCCACAAAGAATG |
| *Asb9* | ACTATAACATCAGCCATC | CCTTGATTCACAGATACT |
| *Best1* | AACTTGAACATTCCAGAG | TCATTAGAGCCTGTATATTG |
| *Prm2* | GGACTATGGGAGGACACA | ATCCTATGTAGCCTCTTACGA |
| *Tnp2* | AAAGTGAGCAAGAGAAAGG | TTGTATCTTCGCCCTGAG |
| **Primers for Genotyping** | | |
| **Gene** | **Forward Primer** | **Reverse Primer** |
| *Lhcgr* wild-type | TGACCTGTTCCTGGGGCTGCTG | AAATGGCCTCAACGGGTGTGCA |
| *Lhcgr* mutant | ATGGGATCGGCCATTGAACAAG | TCAGAAGAACTCGTCAAGAAGGC |
| **Primers for Integration Assay** | | |
| **Gene** | **Forward Primer** | **Reverse Primer** |
| *Lhcgr* | AGCTAATGCCTTTGACAACCTC | CGAGATTAGCGTCGTCCCAT |
| *Cag* | TTCGGCTTCTGGCGTGTGA | GGTGAGAGATAGTCGGGCG |

**Table S3. Primary and secondary antibodies used for immunostaining.**

| **Antibodies** | **Dilution** | **Distributor (Cat.No)** |
| --- | --- | --- |
| Rabbit anti-Pdgfrα | 1:200 | Abcam (ab203491) |
| Rabbit anti-Nestin | 1:200 | GeneTex (GTX133111) |
| Rabbit anti-Ddx4 | 1:400 | CST (8761S) |
| Rabbit anti-Sox9 | 1:200 | Millipore (AB5535) |
| Rabbit anti-Cyp17a1 | 1:200 | CST (94004s) |
| Rabbit anti-Insl3 | 1:200 | Novus (A96525) |
| Mouse anti-Lhcgr | 1:200 | Novus (NBP2-54479) |
| Mouse anti-Tnp2 | 1:200 | Santa Cruz (SC-393843) |
| PNA | 1:400 | Sigma (L7381) |
| Goat Anti-rabbit IgG Alexa 488 | 1:1000 | Invitrogen (A11037) |
| Goat Anti-rabbit IgG Alexa 647 | 1:1000 | Invitrogen (A32733) |
| Goat Anti-mouse IgG Alexa 647 | 1:1000 | Invitrogen (A32728) |

**Video S1. Sperm analysis of the PBS-treated *Lhcgr*^-/-^ mice in pubertal cohort.**

**Video S2. Sperm analysis of the AAV8-Lhcgr-treated *Lhcgr*^-/-^ mice in pubertal cohort.**

**Data file S1. Primary data.**
